# Microbiome diversity of low biomass skin sites is captured by metagenomics but not 16S amplicon sequencing

**DOI:** 10.1101/2025.06.24.661265

**Authors:** Laura Markey, Evan B. Qu, Calen Mendall, Ana Finzel, Arne Materna, Tami D. Lieberman

## Abstract

Established workflows for microbiome analysis work well for high microbial biomass samples, like stool, but often fail to accurately define microbial communities when applied to low microbial biomass samples. Here, we systemically compare microbiome analysis methods —16S rRNA sequencing, shallow metagenomics, and qPCR PMP™ panels—as well as extraction methods across skin swab samples and mock community dilutions. While extraction method minimally impacted results, with no significant signal for method-specific contamination or bias, we observed critical differences in inferred composition across analysis methods for low biomass samples. Metagenomic sequencing and qPCR revealed concordant, diverse microbial communities on low biomass leg skin samples, whereas 16S amplicon sequencing exhibited extreme bias toward the most abundant taxon. Both qPCR and metagenomics showed that female genital tract bacteria dominated the leg skin microbiome in about half of female subjects. Metagenomics also enabled sub-species analysis, which demonstrated that individuals have consistent within-species diversity across high-biomass forehead and low-biomass leg skin sites. This work illustrates that shallow metagenomics provides the necessary sensitivity and taxonomic resolution to characterize species and strain-level diversity in extremely low biomass samples, opening possibilities for microbiome discovery in previously unexplored niches.

**Importance:** Despite the importance of the skin microbiome in health and disease, there have been far fewer studies characterizing the microbiome of skin compared to that of the gut. In part, this is because microbiome methods were initially developed for bacteria-rich samples like stool and these methods perform poorly on bacteria-poor samples like skin swabs. The perceived difficulty of getting reliable data from such low biomass sites has limited the scope and types of analyses performed. Here we demonstrate that shotgun, whole-genome, metagenomic sequencing recovers the full input from even very dilute control community samples, and reveals a highly diverse population even on very low biomass skin sites. We describe a streamlined sample processing and analysis pipeline which can be applied broadly to characterize low biomass microbiome samples and reveal new host-microbiome interactions.

## Introduction

Early studies in the human microbiome field began with sites rich in microbial DNA^1^. The success of these studies generated standardized protocols to robustly profile samples ^2,3^. However, because these approaches were designed for high biomass samples, applying them to very low biomass samples has often led to erroneous results ^4,5^. Low biomass samples are much more susceptible to laboratory contamination and cross-contamination than higher biomass samples in which contaminant DNA is a negligible minority of the total microbial DNA ^5–8^.

The microbial biomass across human skin sites varies dramatically across skin regions with distinct biophysical parameters ^9–11^, with dry skin sites being particularly low biomass and difficult to sample reliably. While there is a significant body of research on the most abundant genera of the skin microbiome which colonize multiple skin environments (including *Cutibacterium*, *Staphylococcus* and *Corynebacterium* ^12–15^), it is possible that underexplored low abundance microbes on dry skin sites also play biologically important roles.

Many aspects of study design can impact the accuracy of inferred species composition, including collection technique, extraction method, and analysis approach. Previous studies comparing collection techniques have shown that swabbing provides the most consistent results for skin microbiome studies ^16–20^ and also results in a higher ratio of microbial to human DNA than other methods ^21–24^. Many studies have compared commercial DNA extraction kits, however, these studies have primarily used high biomass microbiome or mock community samples which are not representative of many skin swab samples ^25–32^.

Published studies to date comparing microbiome analysis approaches produce different expectations for performance when applied to low biomass skin microbiome samples. Both 16S amplicon sequencing and metagenomics have shown to produce equivalent taxonomic abundance for stool samples^33,34^ and high biomass skin sites^35^. However, studies of low biomass environmental samples have reported divergent results, with some favoring 16S^36^ and others favoring metagenomics^37^. In this study, we set out to compare these methods alongside species-specific qPCR panels as an approximate ground truth.

Here we leverage samples with a 10,000-fold range of starting microbial DNA, including high and low biomass skin swabs and a mock community dilution series, to evaluate three methods for DNA extraction and three approaches for DNA analysis. We show that while a qPCR panel, metagenomic and 16S sequencing all perform equally well on high biomass samples, there are significant limitations of 16 amplicon sequencing for recall and diversity recovery in low biomass samples.

## Results

### Collection of skin swabs and construction of microbial load-matched mock communities

The dry skin of the lower leg is colonized by 10^1^-10^4^ colony forming units (CFUs) of bacteria per cm^2^, in contrast with the higher microbial load of oily forehead skin, which can exceed 10^6^ CFUs/cm^2^ ^38–40^. To compare methods across high and low biomass skin samples, we collected replicate skin microbiome samples from the forehead and lower leg of 20 healthy volunteers aged 19-47 using a standardized swabbing protocol (Methods, subject metadata in Table S1). To ensure consistent sampling across DNA extraction methods, a single experimenter collected four replicate swabs from equivalent skin areas (5 cm^2^), in a randomized order across different extraction methods (Fig. 1A, Fig. S1). One sample, collected in 50% glycerol, was stored at −80°C for future analysis of live cells and the remainder were used for DNA extraction (Fig. 1C).

**Fig. 1:**
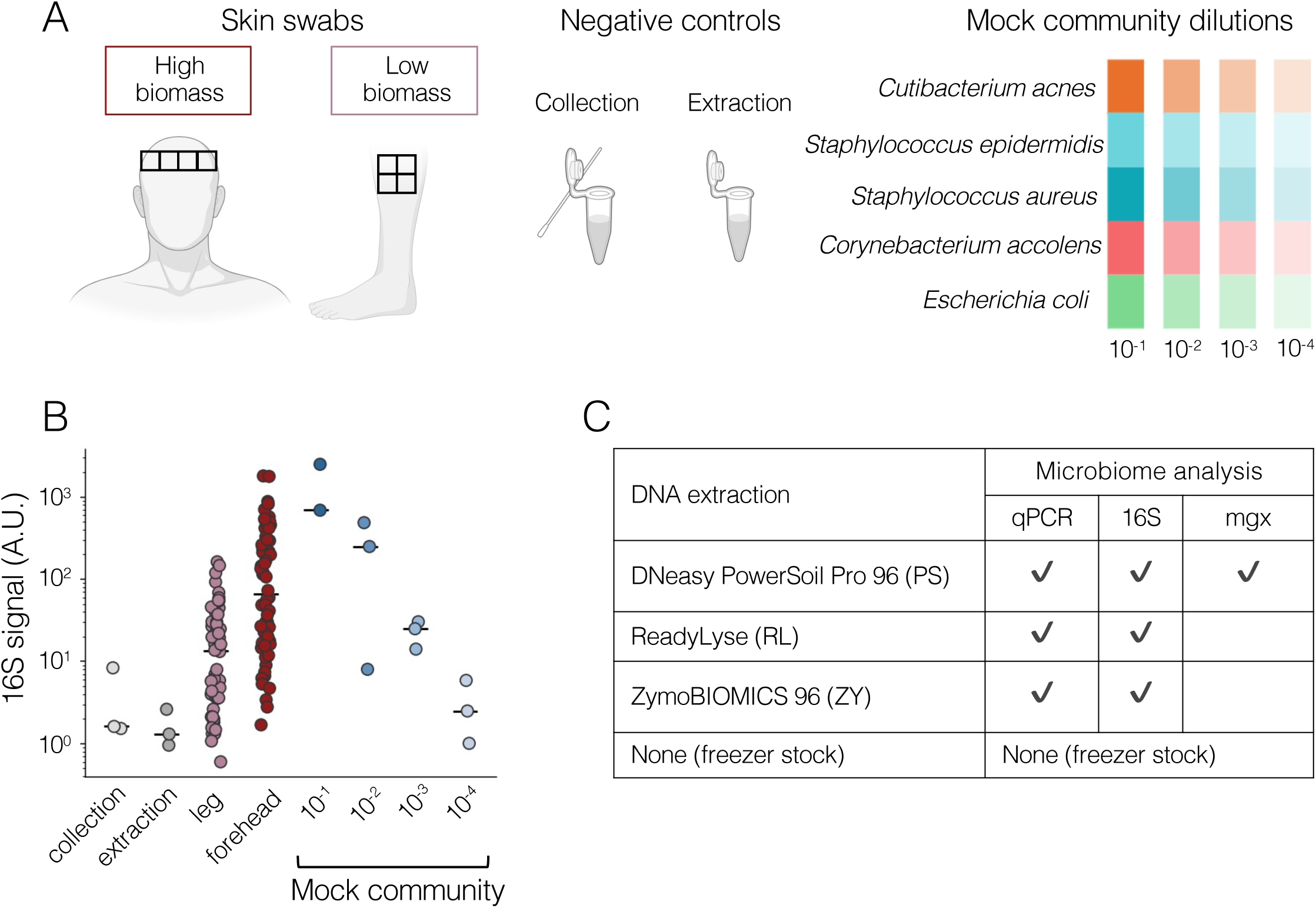
Replicate skin swabs from high and low biomass sites were used to compare DNA extraction and microbiome analysis methods. **A)** Three types of samples were used to assess the impact of DNA extraction and analysis methods on microbiome composition across a range of input bacterial biomass. **Left**: replicate skin swabs from volunteer foreheads (high biomass) and lower leg (low biomass); **Middle**: a “collection” negative control (swab exposed to collection environment) and an “extraction” negative control (reagents added during DNA extraction) were included for each method; **Right**: common skin microbiome species as well as the Gram-negative *Escherichia coli* were combined in equal proportions to form a mock community. **B)** Total microbial DNA was quantified using qPCR and universal 16S (V3 region) primers and normalized to input volume. Each dot indicates a sample. Bars indicate median. Collection N=3, extraction N=3, leg N=58, forehead N=60, mock community N=3 per dilution except 10^-1^ N=2. The 10^-1^ dilution processed with the PS method was contaminated and excluded from analysis. Leg samples identified as likely cross-contaminated excluded from analysis (Fig. S9). **C)** Replicate skin swab samples were randomly assigned to a DNA extraction method or freezer stock; microbiome DNA analysis was performed as indicated in table. Shotgun metagenomics analysis abbreviated as “mgx”.

To assess workflow performance across the biomass range, we created a five-species mock community and profiled a dilution series of this community using all methods (Fig. 1A, N=1 sample per dilution per DNA extraction method). This dilution series spanned 10^5^-10^8^ CFUs/mL, a significantly lower range than commercial mock communities (10^7^-10^10^ CFUs/mL). To confirm that these mock community samples provide a reasonable approximation of skin swab biomass, we measured total bacterial absolute abundance using qPCR with universal primers for 16S rRNA. Indeed, the 16S abundance of all forehead samples and 59/60 leg samples fell within the range of the dilution series (Fig. 1B).

Samples were used to prepare libraries for 16S V1-V3 amplicon and metagenomic sequencing which was performed at MIT by the BioMicroCenter (Methods). Samples were also sent to Bio-Me for analysis with a species-specific qPCR PMP™ panel (Methods). Samples with fewer than a threshold number of reads assigned to microbial taxa were removed from sequencing-based analyses as unlikely to accurately reflect microbiome composition (8 of 120 16S samples; 5 of 40 metagenomic samples; Fig. S2 and Table S2). Samples analyzed by qPCR panel were assessed for bacterial DNA by universal 16S primers; 1/100 samples failed to produce 16S signal.

### Removing likely contaminants using mock communities to identify thresholds

Contamination from the laboratory and cross-contamination from other samples can inflate estimates of bacterial diversity. While previous studies have used a combination of experimental and computational methods to identify likely contaminants, these studies were primarily designed for 16S sequencing data rather than metagenomics. One common approach is to remove all taxa below a single chosen threshold for relative abundance, averaged across experimental samples ^41–43^. However, the appropriate threshold to choose is not always clear.

Another common strategy to remove contaminants is to remove taxa that are at a higher relative abundance in negative control samples than in experimental samples ^44,45^. However, we observed that the most abundant taxa in negative controls were present at similar relative abundances in skin swab samples (Fig. 2A) and included many known residents of human skin. Therefore, a threshold based only on high relative abundance in negative controls would inappropriately remove many true skin microbiome components.

**Fig. 2:**
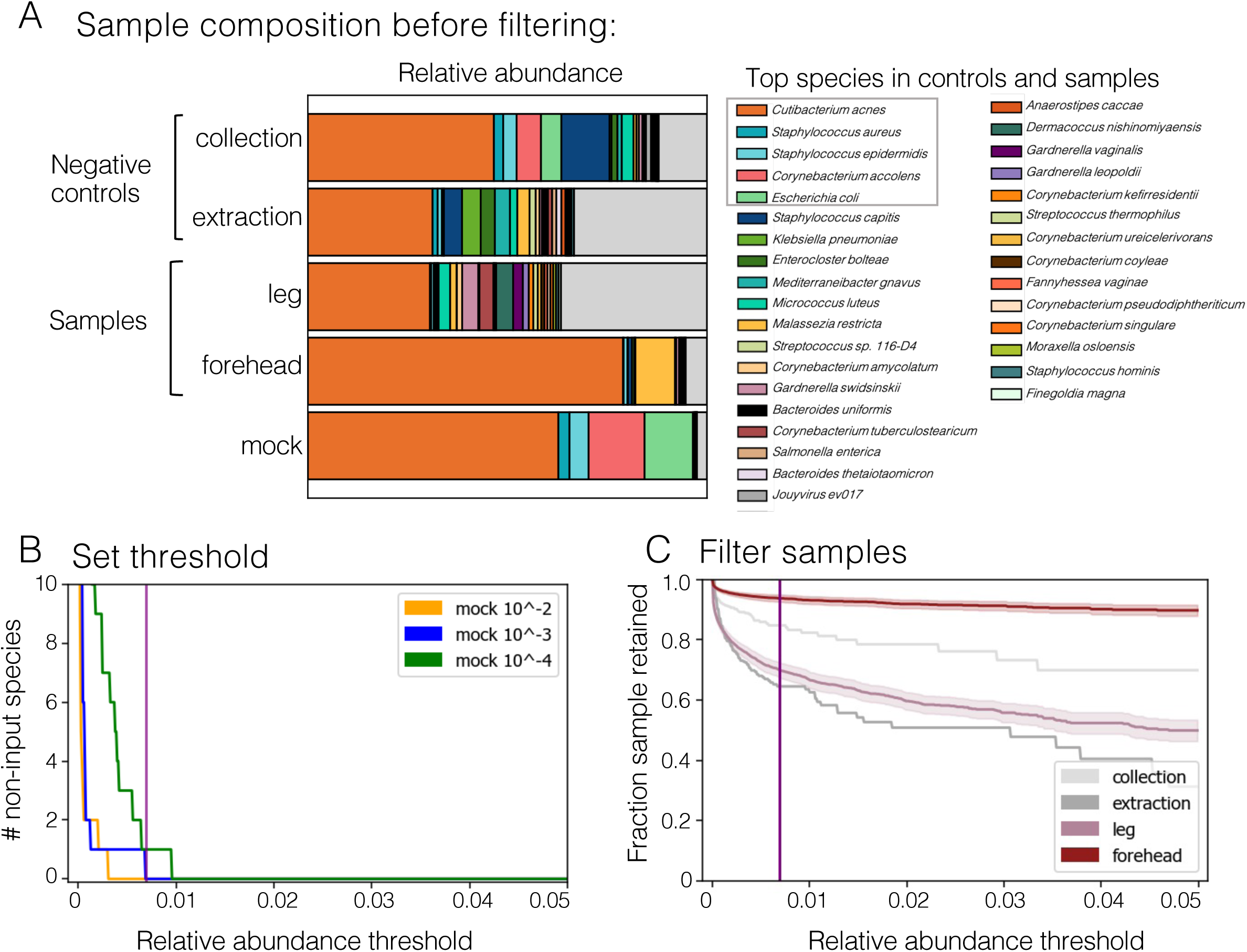
Mock community dilution series is used to set per-sample relative abundance threshold for taxa exclusion. Samples were extracted using the DNeasy PowerSoil Pro 96 kit (PS) and analyzed by metagenomic sequencing. **A)** The average composition of samples in the unfiltered dataset is shown. The 20 most abundant species in the negative control samples as well as the 20 most abundant species in the leg samples are shown in color. **B)** The threshold relative abundance for taxa removal is the value at which the number of non-input species detected in the mock community 10^-3^ dilution is zero (purple line), as taxa present below this threshold are by definition comparable in abundance to contaminants. The 10^-3^ dilution was chosen because the 10^-4^ dilution failed to recall all four input genera for two out of three DNA analysis methods (Fig. S4E). **C)** As the threshold for taxa inclusion increases, the fraction of the original taxonomic composition retained after filtering decreases for low biomass leg samples, which have a higher initial proportion of low abundance species than high biomass forehead samples. Line indicates average; shaded area indicates 95% confidence interval. Leg N=13, forehead N=20, N=1 per collection and extraction controls. The 10^-1^ dilution of the mock community was omitted due to splash contamination during PS DNA extraction. All datasets were individually filtered as shown in Fig. 2B-C; additional datasets shown in Fig. S3. Samples with <100,000 microbial reads excluded from analysis (Fig. S2).

We leveraged a mock community dilution series to set abundance thresholds for taxa exclusion (Fig. 2B), as all non-input species in these samples must originate from contamination or misclassification. For each dataset (unique combination of DNA extraction and DNA analysis approach; 7 datasets total), we chose the abundance threshold that retained all five input species while removing the most non-input taxa (metagenomics shown in Fig. 2B; other datasets Fig. S3) from the 10^-3^ dilution of the mock community. This dilution was chosen in order to standardize contaminant identification across analysis approaches, as the lowest 10^-4^ dilution sample did not contain multiple input genera by 16S sequencing regardless of threshold (Fig. 3A and Fig. S4E). Importantly, this approach enables a taxon to be retained in one sample but removed from others (Methods).

**Fig. 3:**
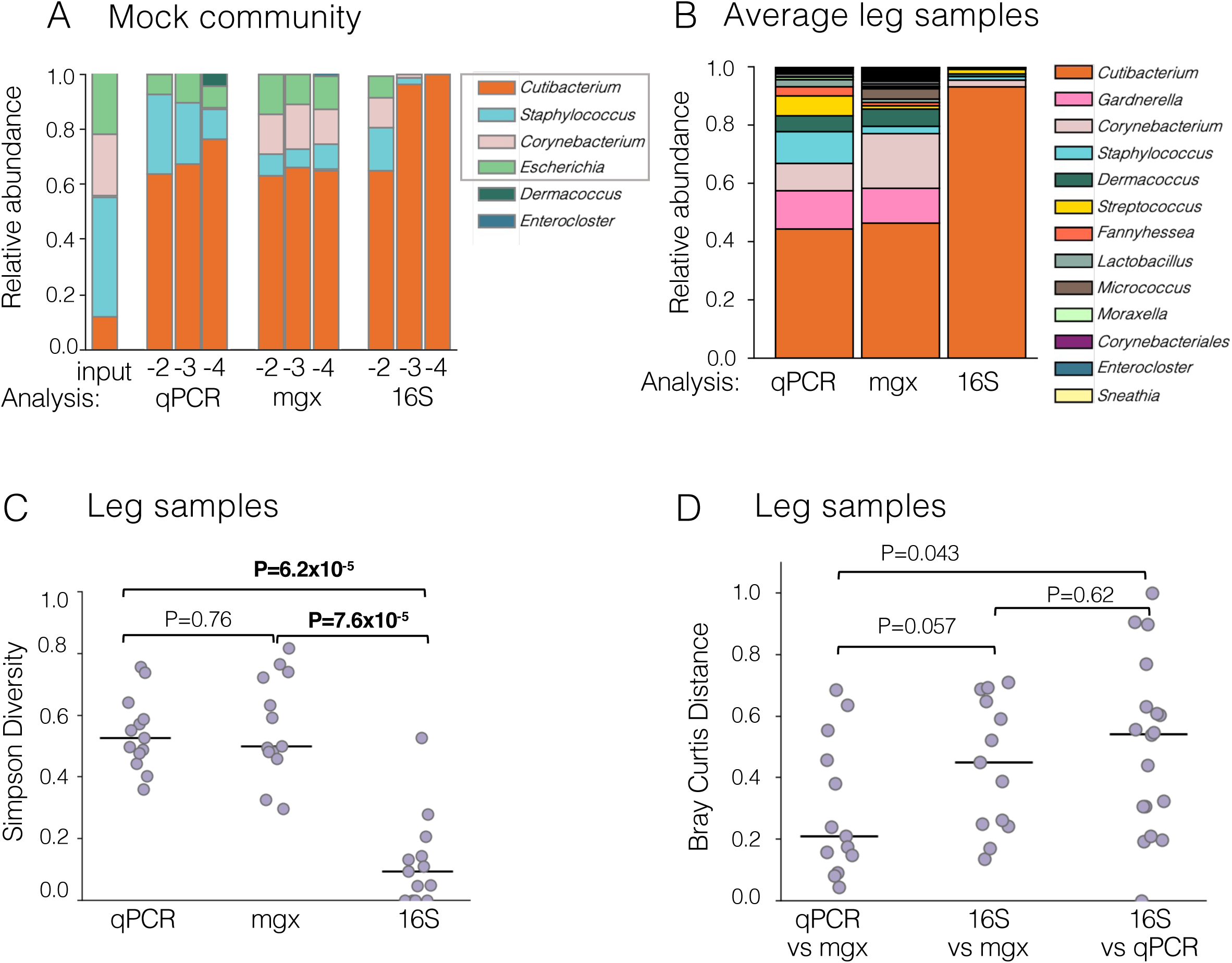
16S sequencing fails to capture diversity of low biomass samples. Samples were extracted using the DNeasy PowerSoil Pro 96 kit and analyzed with a qPCR panel (qPCR), metagenomic sequencing (mgx) and 16S sequencing (16S). Composition is summarized at the genus level. **A)** Mock community samples had a consistent composition across the dilution series when analyzed by qPCR or metagenomic sequencing but 16S sequencing showed an increasing bias towards *Cutibacterium* with dilution. The qPCR panel did not include primers specific for *Corynebacterium accolens*, hence the lack of *Corynebacterium* in the samples. Input genera are shown in a gray box. Each bar presents a single sample. **B)** Both qPCR and metagenomics portray a diverse average leg skin microbiome community while 16S sequencing reflects a *Cutibacterium*-dominant microbiome. All genera present at 10% abundance or higher in at least one sample are shown in a unique color. Bars represent the average of all leg samples included in that DNA analysis method. **C)** Within sample diversity (Simpson metric) was significantly higher when samples were analyzed using qPCR or metagenomics compared to samples analyzed using 16S amplicon sequencing (P=6.2×10^-5^ and P=7.6×10^-5^ respectively). **D)** We observed decreased Bray-Curtis distance between leg samples analyzed using qPCR and metagenomics compared to that between samples analyzed using 16S and metagenomics sequencing (P=0.043). This analysis included only those genera detected by all three DNA analysis methods. Throughout, dots indicate leg swab samples and lines indicate median Significant p-values shown in bold. qPCR N=17, metagenomics N=13 and 16S N=18. Samples which failed internal assay controls (qPCR) or which fell below a microbial read threshold (sequencing assays) not included. Comparable analysis of forehead samples is shown in Fig. S6A-C.

As expected, this filtering approach had the largest impact on the inferred composition of low biomass samples (metagenomics shown in Fig. 2C; other datasets Fig. S3). In the metagenomic data set, this approach removed a median of only 4.6% of the unfiltered taxonomic composition from forehead samples (range 1%-21%) while removing a median 28% from leg samples (range 7.6%-58%). Alternate taxa filtering strategies either fail to remove non-input species from the mock communities or would remove the majority of true species from mock community and skin swab samples (Fig. S5).

### 16S amplicon sequencing underrepresents diversity of low biomass samples

Microbiome analysis methods have distinct advantages for determining taxonomic composition: 16S amplicon sequencing offers reduced cost and database dependencies^46,47^, metagenomics provides higher resolution (including at the sub-species level), and qPCR enables absolute abundance measurements. The set of samples extracted using the DNeasy 96 PowerSoil Pro Kit (PS) method were analyzed using all three methods and the microbiome profiles compared. To fairly evaluate these methods despite their known differences in taxonomic resolution, we analyzed all datasets at the genus level.

We found that 16S sequencing significantly underrepresents microbial diversity in low biomass samples (Fig. 3A-D). While all methods performed comparably with high biomass mock communities, clear differences emerged as samples were diluted. Both metagenomics and qPCR maintained consistent relative abundances of the mock community across the dilution series. In contrast, 16S sequencing showed increasing bias toward *Cutibacterium* with dilution (Fig. 3A). This differential effect of analysis method was mirrored in the analysis of the skin swab samples. High biomass forehead samples showed a limited effect of analysis method – although Simpson diversity was increased in the qPCR dataset compared to either sequencing-based method, the overall composition of forehead samples was not significantly different regardless of analysis methods compared (Fig. S6A-C). Low biomass leg samples showed striking differences as a consequence of analysis method, as both qPCR and metagenomics detected a significantly more diverse microbiome than 16S sequencing (Fig. 3C, P=6.2×10^-5^ and P=7.6×10^^-5^). Additionally, we observed that the overall composition of samples was more similar between qPCR and metagenomics than between qPCR and 16S (Fig. 3D, P=0.043), suggesting that metagenomics more accurately captures the proportions of bacteria in low biomass samples.

We find good correspondence between the relative abundance of 22 genera measured by both qPCR and shotgun metagenomics (Fig. S6D, R^2^=0.90, P=4.2×10^-11^). This strong correlation supports the true sample diversity of the microbiome recovered from lower leg skin, as opposed to misleading diversity from metagenomic classification database errors. The apparent low impact of misclassification is particularly noteworthy because Kraken2/Bracken were used with default parameters against a prebuilt database (based on Refseq genomes) and analysis limited by the 20-80% of reads aligned to human DNA (Fig. S7A). These results demonstrate that a “by-the-book” implementation of shallow metagenomics can yield accurate results for analysis of the skin microbiome.

### Absolute abundance can be estimated from metagenomics for forehead but not leg swabs

Sequencing-based analysis necessarily provides relative abundance measures rather than absolute abundance measures, limiting the interpretability of results. Therefore, there is a great deal of interest in methods to derive absolute abundance information from sequencing data^48–50^. Normalizing to 16S copy number using qPCR to quantify 16S gene abundance is an effective method to approximate absolute abundance^51,52^, however performing qPCR in parallel with sequencing analysis adds to the cost of microbiome research.

Recently, researchers have shown a strong correlation between 16S copy number and the ratio of bacterial to human DNA in gut microbiome metagenomics, thus enabling approximation of total biomass and absolute abundance from metagenomic sequencing alone ^51^. However, gut microbiome samples have significantly higher bacterial biomass than a skin swab sample. We therefore tested this relationship in our data and observed a strong correlation (R^2^=0.3, P=0.00098, Fig. S7C) between the microbial to human DNA ratio and 16S copy number when all skin samples were analyzed together and in higher biomass forehead samples alone (R^2^=0.51, P=0.00044, Fig. S7C). Interestingly, no correlation between the two metrics was found in low biomass leg skin samples (R^2^=0.0018, P=0.89, Fig. S7C). Our results support the findings of Tang et al^51^, and show that for microbiome samples with sufficient biomass the human-to-microbe read ratio is an appropriate metric for absolute abundance estimation.

### Kit-based methods outperform direct enzyme approach for low biomass samples and introduce minimal contamination

We compared three DNA extraction approaches: DNeasy 96 PowerSoil Pro (PS), ZymoBIOMICS 96 (ZY), and direct ReadyLyse addition (RL) across sample replicates from the same individuals. For RL, enzyme was added directly to collection buffer and the lysis product was diluted 1:1 with UltraPure water and then used directly for PCR; for the other two methods manufacturer’s recommendations were followed, including both lysis and on-column purification (Methods). Each sample was analyzed using universal 16S qPCR, 16S sequencing and a species-specific qPCR panel.

Using universal 16S qPCR to quantify the total bacterial DNA per sample, we observed that kit-based methods (PS and ZY) resulted in significantly higher DNA yields than RL from high biomass forehead samples (Fig. 4A, P=8.5×10^-6^ and P=5.9×10^-6^). We found similar results for lower leg samples (Fig. 4A, P=0.0075), though there was not a statistical difference between ZY and RL for these lower biomass samples. In accordance with this observation of low yields, five samples extracted via RL and three samples via ZY failed to meet our per-sample 16S read threshold, while all samples extracted via PS met this threshold (Fig. S2B, Table S2). In addition, PS and ZY more accurately recovered the complete mock community from dilute mock community samples compared to RL (Fig. S4E). Together, these results suggest that mechanical and chemical lysis methods like PS and ZY are most appropriate for low biomass applications, with a slight advantage for PS.

**Fig. 4:**
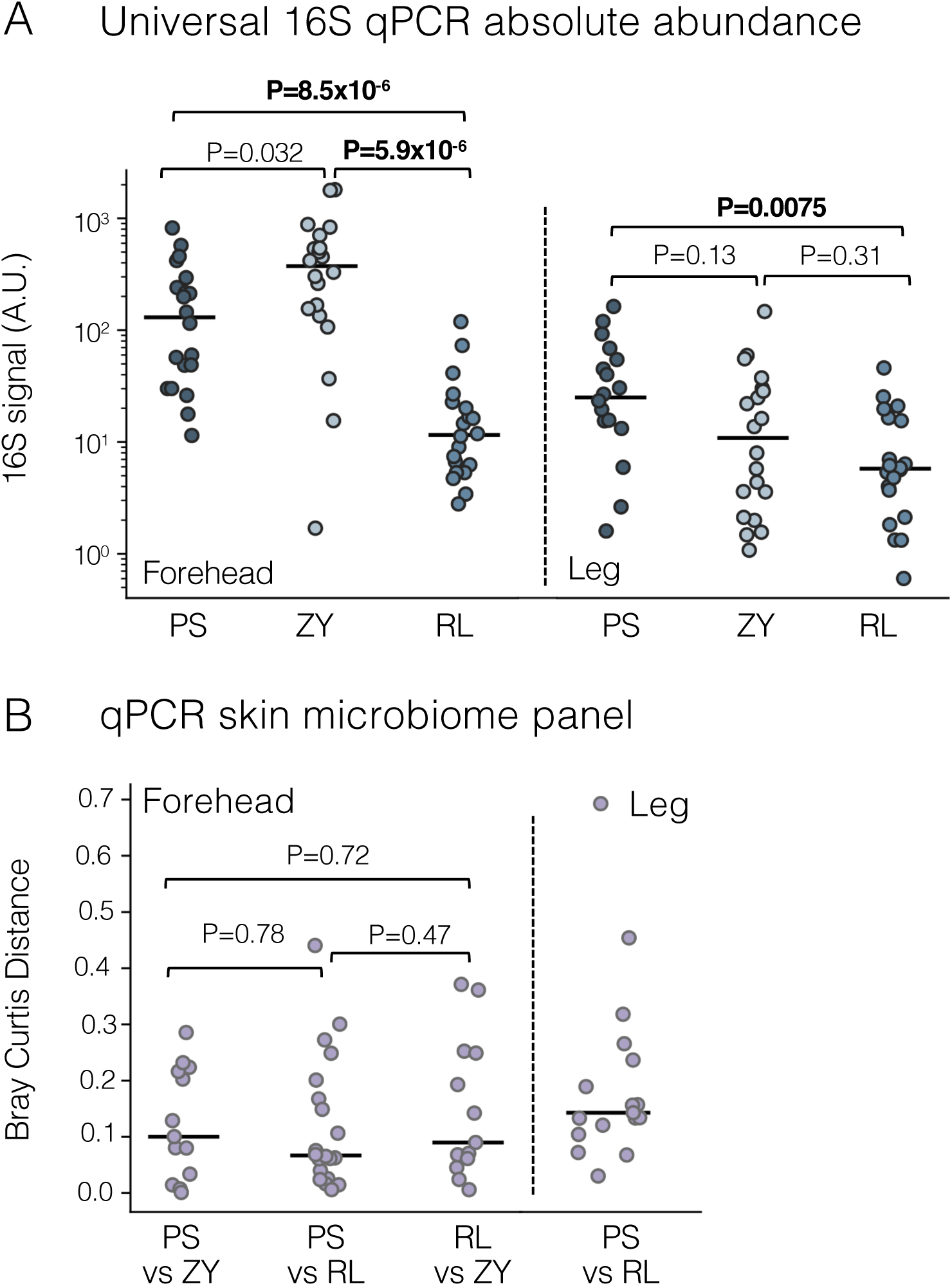
DNA extraction via mechanical and chemical lysis increases yield and does not introduce major contaminants. Three DNA extraction methods were used to process replicate forehead and leg skin swab samples: DNeasy PowerSoil 96 kit (PS), direct lysis using ReadyLyse enzyme (RL), and ZymoBIOMICS 96 DNA kit (ZY). **A)** Total bacterial DNA yield per swab differed significantly between extraction methods for both forehead and leg samples (P<0.008). 16S signal was assessed using universal 16S qPCR and adjusted for input volume. N=20 for all sites and methods except PS leg N=18**. B)** Analysis by qPCR skin microbiome panel shows that despite differences in yield, there is no difference in Bray-Curtis distance between pairs of samples extracted using kit-based methods (PS and ZY) compared to pairs of samples including a kit-based method and RL (P>0.40). Forehead: PS N=20, RL N=20, ZY N=13; leg PS N=17, RL N=20. The comparable composition further supports our finding that kit reagents are not a major source of contaminants (Fig. S8). The qPCR analysis of leg swab samples does not include ZY leg samples due to evaporation of ZY samples prior to qPCR panel submission. Bray-Curtis distance between sample replicates extracted by different methods analyzed by 16S sequencing shown in Fig. S4F. Significant p-values shown in bold.

Despite extraction-dependent differences in DNA yield, DNA extraction method had minimal impact on skin microbiome profiles by either qPCR panel (Fig. 4B and Fig. S4) or 16S sequencing (Fig. S4) for the samples that passed filters. While we were expecting to find contaminants specific to each extraction method, no genus was significantly associated with any extraction method (Fig. S8, Table S3). Notably, negative control samples from PS had significantly higher absolute abundance as assessed by qPCR than RL and ZY (Fig. S8A); PS samples were lysed in 96-well format in contrast to RL and ZY, which used within-tube lysis, potentially implicating well-to-well transfer as the primary source of contamination.

### Lower leg skin microbiome reflects higher biomass microbiome sites

Using our optimized low biomass protocol (PS extraction followed by metagenomic sequencing), we characterized the dry skin microbiome of the lower leg. Despite extremely low biomass (median 0.023 ng/μl total DNA concentration, Table S1), we successfully analyzed 13/20 samples after excluding five due to insufficient microbial reads and two due to well-to-well contamination (Fig. S2A and Fig. S9B-C).

Leg samples were distinct from negative controls both in total biomass (Fig. 1B, P=0.0059) and in overall composition (Fig. S9A, PERMANOVA P=0.15). Leg samples had high species and genus-level diversity within and across subjects (Fig. 5A). We detected both well-known skin commensals (*Corynebacterium* and *Staphylococcus*), as well as less well-studied genera including *Dermacococcus* and *Micrococcus*.

**Fig. 5:**
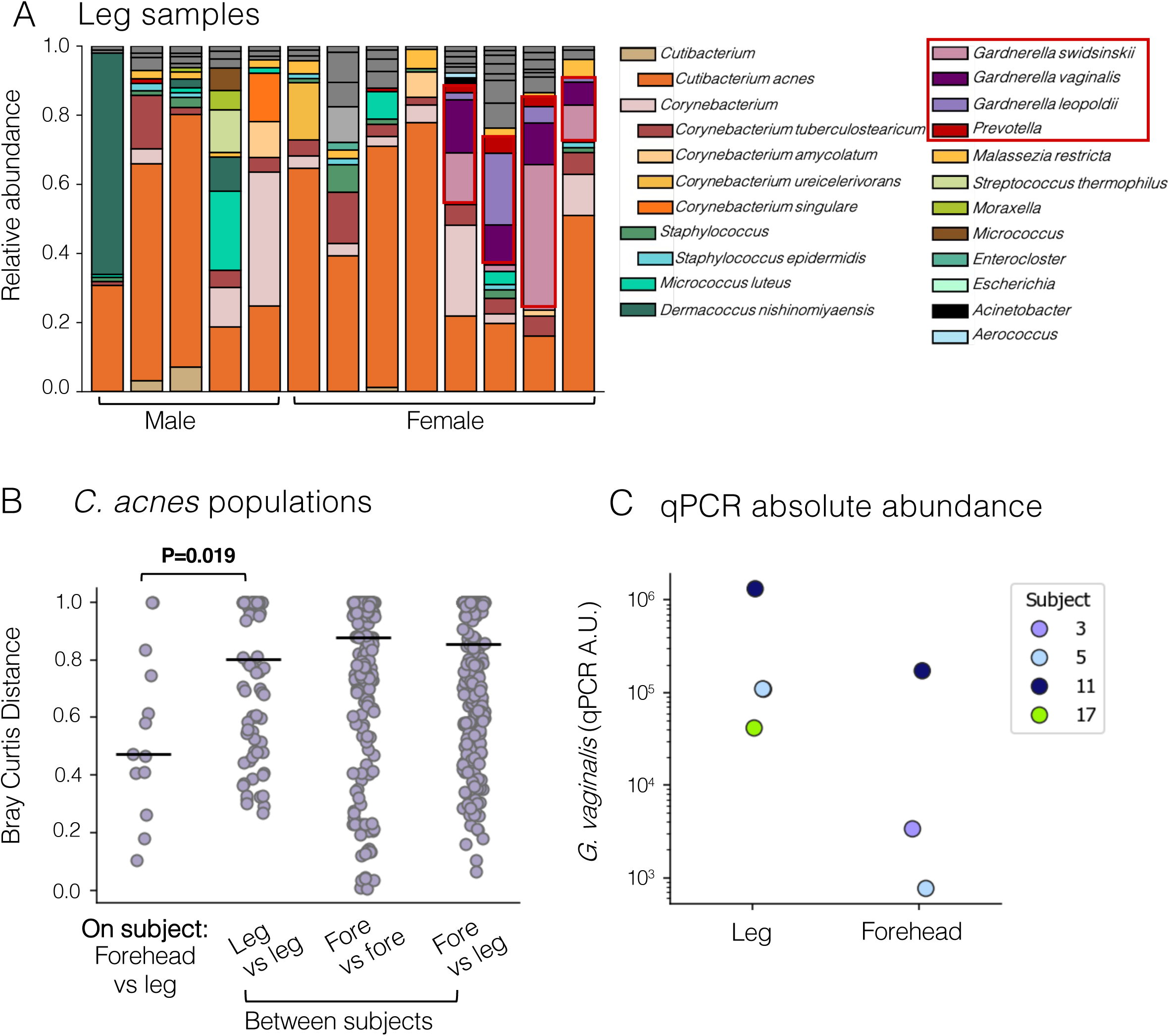
Diverse leg skin microbiome shows transfer from other microbiome body sites. DNeasy PowerSoil 96 samples were analyzed using metagenomic sequencing and the leg skin microbiome profiled at the strain, species and genus level. **A)** The leg skin microbiome is diverse within and across individuals -- in four out of eight female subjects, it contains a substantial proportion of female genital tract (FGT) microbiome species presumably transferred from the FGT. Samples are ordered by subject sex, with the subset of female subjects whose leg skin microbiome contained >20% common FGT species at the far right. Species and genera with an average abundance >10% in at least one sample are shown with a unique color. Red boxes indicate FGT taxa. **B)** Sub-species relative abundance analysis (PHLAME) of *C. acnes* shows that forehead and leg *C. acnes* populations on the same subject are more similar than *C. acnes* populations across all leg samples (P=0.019). The Bray-Curtis distance between samples was calculated within and between subjects and within and between body sites. Dots indicate pairs of samples and lines indicate median. Leg N=13 and forehead N=20. **C)** A FGT qPCR panel validated the presence of FGT microbe *Gardnerella vaginalis* in the leg samples of four female subjects and also detected *G. vaginalis* in the paired forehead samples of three out of the four subjects, supporting transfer of FGT bacteria across skin sites. Dots represent samples. Leg N=17 and forehead N=20. Forehead microbiome profiles shown in Fig. S10.

In addition, we discovered that female genital tract (FGT) microbes made up a substantial proportion of the leg microbiome in four out of eight female subjects, predominantly *Gardnerella* and *Prevotella* species (Fig. 5A). This unexpected finding was validated by FGT qPCR panel, confirming increased absolute abundance of *Gardnerella vaginalis* in all four leg samples and three of four forehead samples from these subjects (Fig. 5C, Fig. S10B). We exclusively observed FGT microbe-dominated leg skin samples in female subjects, suggesting direct transfer of microbes between the FGT and skin microbiome.

Lastly, we demonstrated the utility of metagenomics for intra-species analysis by focusing on *Cutibacterium acnes.* Using PHLAME^53^, which classifies intraspecies diversity at conserved evolutionary groupings, we find that individuals maintain consistent *C. acnes* populations across body sites; each subject’s forehead and leg harbor highly similar clade proportions distinct from other subjects (Fig. 5B and Fig. S10D). No clade of *C. acnes* showed skin-site specific enrichment (Fig. S10C). This result extends previous findings of strain sharing across other skin sites^11^ to include the lower leg.

## Discussion

In this study we perform a comprehensive comparison of microbiome analysis methods and conclude that for low biomass samples, metagenomic sequencing is critical for accurate results. As sample biomass decreases, 16S sequencing becomes heavily biased towards dominant taxa, while shotgun metagenomics and qPCR effectively recover microbiome diversity across the sample biomass spectrum, with minimal loss of mock recall or sample diversity (Fig. 3).

As the cost of short-read sequencing continues to decrease with multiple competitors in the space, there is no longer a significant difference in sequencing cost between 16S amplicon sequencing and shotgun metagenomics (∼$15 materials cost per sample when pooling 100 samples at our sequencing facility). We further reduced the cost of metagenomics by using a modified version of the Hackflex protocol^54^ to prepare libraries.

In contrast with previous studies^16,28,55,56^, we did not observe a strong signal for reagent-derived contamination. Instead, contaminant DNA primarily stemmed from well-to-well sample transfer, with the highest rate of contamination observed in the plate-based PS lysis rack (Fig. S8 and Fig. S9). We recommend within-tube lysis (rather than within-plate lysis) to limit opportunities for such contamination and other process controls in line with previous studies^57^.

While these tube-based kits can add to the cost of microbiome studies (∼$5-10 materials/sample), our results demonstrate that the combination of mechanical lysis, chemical lysis, and column purification significantly improves both mock community recall and human sample recovery (Fig. 4 and Fig. S4E), justifying the investment.

Whether qPCR or metagenomics is more appropriate for assaying low biomass samples depends on the specific goals of a given study. Advantages of qPCR include extremely sensitive detection of bacterial DNA unaffected by host DNA and quantification of absolute abundance unlimited by biomass – however its utility for exploratory microbiome research is limited by the inherently targeted nature of the assay. Our initial qPCR analysis misrepresented the microbiome of a subset of leg skin samples dominated by female genital tract (FGT) microbes not included in the skin-specific qPCR panel. Based on this metagenomics data, we then added the PMP^TM^ vaginal microbiome qPCR panel to our qPCR analysis, producing results that correlated well with the metagenomics data (Fig. 5 and Fig. S10B).

While we were not surprised to observe a diverse microbiome on lower leg samples, based on other findings at dry skin sites^11,58^, an unexpected finding was that the lower legs of 50% of female subjects in our study were heavily colonized by *Gardnerella sp*, typically found in the FGT. This genus is comprised of facultative anaerobes which could plausibly persist once transferred to leg skin. Previous studies have found *Gardnerella vaginalis* on the moist skin of the elbow crease,^11,21^ finger webbing and thigh crease^11^ as well as in trace abundance on the dry skin of the palm and forearm^11^, but we are unaware of other reports of its dominance of a dry site. Previous studies of the vaginal microbiome in healthy, industrialized Western cohorts generally report a *Lactobacillus*, rather than *Gardnerella,* dominant vaginal microbiome^59,60^. While *Lactobacillus* was also present in leg skin samples, its abundance was much lower (Fig. S10B). Studying the basis of *Gardnerella’s* relatively high abundance on the lower leg and the contribution of transferred microbes to the active skin microbiome more generally is an important area for further study.

Altogether this work emphasizes the increased sensitivity achievable from metagenomics over amplicon-based approaches^37,61^ and provides a detailed roadmap for studying low biomass skin sites.

## Methods

### Study design and sample collection

20 healthy volunteers were recruited and samples collected through the Center for Clinical and Translational Research (CCTR) at Massachusetts Institute of Technology (MIT). All study procedures were approved by the MIT Institutional Review Board (IRB # 2102000324) before data collection began. This study included 14 female and 6 male subjects ranging in age 19 to 47 (full metadata in Table S1). First, the forehead was washed using a Cetaphil cleansing wipe followed by a gentle saline wash. While the forehead dried, the sample collector collected leg swab samples as follows. A sterile plastic template was attached to the subject’s lower leg and a Puritan foam swab was pre-wetted in TESX buffer (10mM Tris-HCl, 1mM EDTA, 100mM NaCl, 1% vol/vol Triton-X100) and then rubbed firmly against the skin within the quadrant A square 40 times using both sides of the swab. The swab was then placed in a collection tube according to DNA extraction protocol specific parameters. This process was repeated three times for a total of four replicates from the leg. Then the sample collector changed gloves and placed a sterile template on the forehead and repeated the swabbing protocol for each of the four quadrants on the forehead. After all samples were collected, they were placed in liquid nitrogen to freeze and then kept frozen on dry ice until transfer to −80°C freezer.

### Mock community sample generation

Bacterial species were individually grown in liquid culture then combined in equal proportions and cryopreserved by adding 10% DMSO (vol/vol). Mock communities were divided into 100ul aliquots and frozen at −80°C until used for DNA extraction. Colony-forming-unit (CFU) plating of individual cultures was used to confirm the proportions of CFUs used for mock community. CFU plating results reported as “input” in Fig. 3A reflect a much lower proportion of *Cutibacterium acnes* than is reflected in DNA analysis results; we believe this is likely because the slow growing *C. acnes* cultures were in exponential phase when harvested after 18 hours of growth while the more rapidly growing *Staphylococcus*, *Escherichia coli* and *Corynebacterium accolens* cultures were likely in stationary phase at 18 hours of growth. This would result in more DNA per cell in the *C. acnes* cultures than in the other species.

*Cutibacterium acnes* (lab strain collection) was grown in RCM media at 33°C under anaerobic conditions for 72 hours and then diluted 1:100 and grown overnight. *Staphylococcus aureus* USA300 LAC (gift from Prof. Isaac Chiu) and *Staphylococcus epidermidis* (lab strain collection) were cultured in TSB at 37°C with shaking for 18 hours. *Escherichia coli* MG1655 (gift from Prof. Gene-Wei Li) was cultured in LB at 37°C with shaking for 18 hours. *Corynebacterium accolens* (ATCC collection, #49725) was cultured in BHI 0.1% Tween80 (vol/vol) at 37°C with shaking for 18 hours.

Prior to DNA extraction, a mock community aliquot was thawed and vortexed and diluted using PBS. Diluted mock community samples were then immediately used for DNA extraction.

### Microbiome sample DNA extraction

For all DNA extraction methods, samples were thawed and handled in a biosafety cabinet for all steps preceding sample lysis. All post-lysis steps of the DNA extraction and library preparation protocols were carried out in a lateral flow hood to limit transfer of DNA between wells.

#### DNeasy 96 PowerSoil Pro Kit (PS)

Sample DNA extraction was carried out as per kit manual with some modifications as noted. Briefly, skin sample swab heads and 250μl of collection buffer (10mM Tris-HCl, 1mM EDTA, 100mM NaCl) were transferred to a well of the PowerBead Pro plate and 800μl of Solution CD1 was added per well and the plate was sealed using sealing film. For laboratory control samples, 250μl of collection buffer plus the negative control swab or 250μl of mock community dilution were added to the well along with 800μl of Solution CD1. Samples were homogenized using the TissueLyser with two rounds of shaking for 5 minutes at 25 Hz. Kit reagents were used to remove inhibitors and wash DNA as per protocol and DNA was eluted from kit columns using 50uμl of Solution C6 and stored at −20°C.

The 10^-1^ dilution of the mock community was contaminated during transfer to the PowerBead Pro plate and thus excluded from analysis (across DNA analysis methods).

#### ZymoBIOMICS 96 DNA Kit

Skin swab samples were collected directly into lysis tubes (containing 1ml DNA/RNA Shield and microbial lysis beads). 250μl of negative control sample or mock community dilution was transferred to a lysis tubes. DNA extraction was carried out as per kit manual with the following modifications. Tubes were homogenized using the TissueLyser with two rounds of shaking for 5 minutes at 30 Hz. Tubes were then centrifuged at 12,000xg for 1 minute to pellet lysis beads and 400μl of lysate was transferred to a 2mL 96-well plate. 1200μl of ZymoBIOMICS DNA binding buffer was added to the sample lysate and mixed by pipetting. 800μl of this mixture was transferred to the DNA binding column of the Zymo-Spin I-96-Z plate and samples were washed as per protocol using kit reagents. DNA was eluted from kit columns using 20μl of UltraPure water and stored at −20°C.

Samples extracted using this method showed clear signs of evaporation after multiple months of storage at −20°C. 16S sequencing and universal 16S qPCR analysis were performed at MIT within one month of DNA extraction and thus not impacted by evaporation but qPCR panel analysis was performed several months later and all leg samples and 6/20 forehead samples were eliminated from analysis due to uninterpretable results from low remaining sample volume.

#### ReadyLyse: Direct enzymatic lysis

100μl of negative control sample or mock community dilution was used for DNA extraction. Samples were collected into 0.5mL tubes containing 100μl of collection buffer (10mM Tris-HCl, 1mM EDTA, 100mM NaCl). 4μl of ReadyLyse (1250 units/μl) was added directly to each 100μl sample. Samples were incubated at room temperature for 19 hours.

Samples were then diluted two-fold in UltraPure water prior to PCR amplification or library preparation and stored at −20°C. Remaining lysate was frozen at −80°C.

### 16S amplicon sample preparation, sequencing and analysis

The 16S region of the bacterial rRNA gene was amplified using V1-V3 primers (27F - 543R). Libraries were prepared for sequencing following the Hackflex protocol^54^. Nextera-compatible primers (IDT) were used for index PCR and amplicons were purified using DNA-binding beads (Cytiva) for size selection. Nextera libraries were converted for Element sequencing using the Element Cloudbreak Freestyle conversion chemistry and then sequenced on an Element AVITI using 150bp paired-ends reads to an average depth of 50,000 reads per sample. All data processing was done using QIIME2 (v2021.2)^62^. Only forward reads were used, as reverse read quality was too low to overlap pairs. Adapters were trimmed using the cutadapt plugin for QIIME2, and data were denoised using DADA2^63^to generate the amplicon sequence variant (ASV) table. A custom classifier based on the SILVA database (v132) ^64^ was used for taxonomic assignment. Exported taxa abundance was analyzed and visualized using custom Python scripts. Taxa unassigned at the phylum level and below, as well as taxa assigned to eukaryotes, were removed and abundance renormalized. Because *Finegoldia magna* abundance was highly discordant across DNA analysis methods, ASVs classified as *F. magna* were removed and data renormalized. Samples with fewer than 500 reads assigned to bacterial taxa were removed from analysis (ReadyLyse: 3/20 leg samples and 2/20 forehead samples; ZymoBIOMICS 3/20 leg samples. No PowerSoil samples were removed; Fig. S2). Sample sequencing statistics included in Table S2. Analysis was performed using genus-level taxonomic classification. Further taxa filtering is described below.

### Shotgun metagenomic sample preparation, sequencing and analysis

PowerSoil samples were used for shotgun metagenomic sequencing. Libraries were prepared for sequencing following the Hackflex protocol^54^. Nextera-compatible primers (IDT) were used for index PCR and DNA fragments were purified using DNA-binding beads (Cytiva) for size selection. Nextera libraries were converted for Element sequencing using the Element Cloudbreak Freestyle conversion chemistry and then sequenced on an Element AVITI using 150bp paired-ends reads to an average depth of 3,000,000 reads per sample by the MIT BioMicro Center.

Adapters were removed from metagenomic sequencing reads using cutadapt (v.1.18) and reads were filtered using sickle (v.1.33; -g -q 15 -l 50 -x -n). Reads were aligned to the human genome to discard host DNA sequences; only unaligned reads were used for further analysis. Kraken2 was used to classify metagenomic reads to the species taxonomic level using the RefSeq reference database and taxonomy; bracken was used to estimate relative abundance of bacterial species per sample. Reads assigned as eukaryotes were removed and data renormalized. Because *Finegoldia magna* abundance was highly discordant across DNA analysis methods, reads which aligned to *F. magna* were removed from shotgun metagenomics and data renormalized. Samples with fewer than 100,000 reads assigned to microbial reference genomes were eliminated from analysis (5/20 leg samples and 0/20 forehead samples removed, Fig. S2). Sample sequencing statistics included in Table S2. Further taxa filtering is described below.

### Quantitative microbiome PCR

The PMP™ Comprehensive Skin Microbiome panel v2.0 (Bio-Me, Oslo, Norway) was used in this study. The Precision Microbiome Profiling (PMP™) panel is comprised of TaqMan^™^ qPCR assays for 48 bacterial species and subspecies and 5 *Malassezia* species, a *Malassezia* genus-specific assay, a universal assay targeting bacterial 16S rRNA genes, and an internal positive control (Table S10). Bio-Me carries out the in silico design of PMP™ assays using proprietary software and databases to identify suitable genomic regions for TaqMan™ assays – consisting of a fluorescently labeled probe and two PCR primers – for each microbial taxon of interest. PMP™ primer pairs and probes were custom-manufactured, and preloaded onto TaqMan^™^ OpenArray^™^ plates by Thermo Fisher Scientific (Waltham, US).

TaqMan^™^ Universal DNA Spike-In Control was added to TaqMan^™^ OpenArray^™^ Genotyping Master Mix (1:6) (Thermo Fisher Scientific, Waltham, US). 3 µL of the resulting mix and 2 µL of sample DNA were transferred to a well of a 384-well plate. The 384-well plate was spun at 687xg for two minutes and placed into the OpenArray™ AccuFill™ System (Thermo Fisher Scientific, Waltham, US), to load the samples into the custom-manufactured PMP^™^ qPCR panels. The next steps were carried out according to the manufactureŕs instructions. Panels were analyzed by the QuantStudio™ 12K Flex Real-Time PCR System following the default TaqMan^™^ OpenArray^™^ Real-Time PCR Plates Protocol (Thermo Fisher Scientific, Waltham, US). The work was performed at Bio-Més laboratory (Oslo, Norway). Standard curves, which are routinely generated for the in vitro performance evaluation of individual assays were used to convert the quantification cycle (Cq) value into absolute quantities for each target taxon (reported as number of genomic copies per µL).

Absolute abundance of the species *Finegoldia magna* was discordant with shotgun metagenomics abundance data using both raw read-mapping to the *F. magna* genome with bowtie2 and relative abundance of *F. magna* as classified by kraken2/bracken therefore *F. magna* was removed from all datasets and remaining taxa renormalized.

### Computational removal of likely contaminant DNA signal

#### Threshold abundance method developed for this paper

DNA signal was classified as indistinguishable from contamination when it was detected at or below an abundance threshold. This abundance threshold was set individually for each dataset (unique combination of DNA extraction method and DNA analysis method) in order to capture batch differences in degree and identity of possible false-positive DNA. We used the mock community dilution series to choose this threshold as the mock community samples have a known composition of 5 input species. For each dataset, we chose the lowest abundance threshold that retained all 5 input species and removed the most non-input taxa (Fig. 2B). For 16S and shotgun metagenomic sequencing this number was a relative abundance value while qPCR panel data was filtered using the absolute abundance values.

Values present in a dataset below the abundance threshold were changed to 0 to omit that potentially false-positive signal from confounding the analysis. Data was then renormalized to 1 for relative abundance sequencing datasets. For the qPCR panel data below the threshold was changed to 0 and no other change was required. Taxa that were now <0 across all samples were omitted from analysis.

#### Alternative methods for false-positive DNA sequence identification and removal

We compared our taxa filtering strategy to three other commonly used methods for false-positive DNA removal carried out on the shotgun metagenomics dataset as follows (Fig. S5). The “top 10 blank” method removes DNA based on its presence in negative control samples. The 10 species with the highest average relative abundance across the two negative control samples were removed from all samples and the data renormalized to 1. The “1% threshold” method removes all species with an average relative abundance equal to or less than 1% from the dataset across all samples. Data is then renormalized to 1. The “decontam” method refers to the R software package “decontam” which uses the abundance of a species in sample and negative controls as well as the DNA concentration of those samples to identify likely contaminants. We used decontam v1.22.0 (dependencies ggplot2 v3.5.1, reshape2 v1.4.4, stats v4.3.2) and the default settings (p-value associated with accurate contaminant identification of 0.1) to generate a list of contaminants both from the “frequency” contamination identification mode and the “prevalence” contamination identification mode. Species identified as possible contaminants using either mode were removed from the dataset and data renormalized.

### Identification and removal of samples with splash-over DNA signal from neighboring wells

We analyzed the shotgun metagenomics dataset to identify well-to-well contamination in the DNeasy 96 PowerSoil samples and the qPCR dataset to identify well-to-well contamination in the ZymoBIOMICS and ReadyLyse datasets.

To identify possible well-to-well contamination, we calculated the Bray-Curtis distance between samples from different subjects as well as the number of species per sample. Pairs of samples with a small Bray-Curtis distance and more than the median number of species present were called as possibly contaminated (Fig. S9B). We then compared the composition of these suspect sample pairs across methods – samples which were highly similar by one method but not similar by other methods were removed (Fig. S9C). We removed from analysis the sample from the pair which had lower biomass as calculated by 16S qPCR using universal primers. This method removed two leg swab samples from the PowerSoil 96 sample set and no samples from the ZymoBIOMICS or Readylyse sample sets.

### Sub-species analysis for *Cutibacterium acnes*

Strain-level analysis of *C. acnes* populations in skin swab samples was performed using PHLAME, a tool which uses bacterial species phylogeny to infer the sub-species identity of *C. acnes* reads and generate a phylogenetic abundance profile of *C. acnes* strains based on metagenomic data, described in detail in Qu et al, 2025^53^. Briefly, the shotgun metagenomic dataset was aligned to the *C. acnes* genome using bowtie2 and single-nucleotide variants (alleles) from samples were compared to those present in a *C. acnes* phylogenetic tree in order to classify metagenomic abundance data at the sub-species level. This strain-level relative abundance profile was used to compare the *C. acnes* populations present within and between subjects.

### Universal 16S qPCR sample analysis

All samples were diluted 10^-1^ in UltraPure water; this was used for the template for the qPCR reaction. We used primers specific for the 16S V3 region expected to universally amplify bacterial rRNA^65^. 20μl reactions were set up as follows: 10μL of 2X SYBR MasterMix, 0.5μL F primer (10μM), 0.5μl R primer (10μM) and 7.4μL of UltraPure water. We used a Lightcyler 400 to conduct qPCR reaction and converted the Ct values to arbitrary units using the relative abundance method and comparing sample Ct values to those from a dilution series of mock community samples. Arbitrary units 16S units per sample were then adjusted to estimate the biomass of the starting swab sample based on differences in the volume of swab sample used as input to the DNA extraction method: PowerSoil 96 samples were collected into 1000μL of buffer and 250μL was used for DNA extraction (1000/250=4) therefore values were multiplied by 4; ReadyLyse samples were collected into 100μL of buffer, all of which was used for digestion, and then diluted 2-fold prior to analysis therefore values were multiplied by 2; ZymoBIOMICS samples were collected into 1000μL of buffer and 400μL were used for DNA extraction (1000/400=2.5) therefore values were multiplied by 2.5. These adjusted 16S values in arbitrary units are shown in Fig. 1B.

## Statistics

All statistical analysis was performed using scipy (v1.14.1). Pairs of experimental groups were compared using the Mann-Whitney U test. When applicable, the Bonferroni correction was used to adjust the p-value significance cut-off for multiple hypothesis testing and an alpha of 0.05.

## Supporting information

Table S4-S9 abundance tables

## Data availability

Sequencing files are available to download from the NCBI SRA database BioProject ID PRJNA1277168.

Sample and subject metadata are described in Supplemental Table 1. Statistics describing sequencing files are available in Supplemental Table 2. Raw and filtered abundance tables for 16S, metagenomic and qPCR panel analysis are available as Supplemental Tables 4-9. The *C. acnes* phylogroup relative abundance output from PHLAME is available as Supplemental Table 10. The list of species included in the Bio-Me PMP^TM^ qPCR panels is included as Supplemental Table 11 and 12.

Code used to analyze and visualize data starting from raw abundance tables is available on Github at https://github.com/lmarkey/lowbiomass_microbiome

## Acknowledgements

We would like to acknowledge the work, advice and expertise of Stuart Levine and all of the staff members of the MIT BioMicroCenter and thank them for their assistance in performing 16S and metagenomic sequencing. We would like to acknowledge the MIT Office of Research and Computing for providing high performance computing resources. We would also like to acknowledge staff members of the MIT Center for Clinical and Translational Research (CCTR), particularly Tatiana Levkovich Urman (Senior Clinical Research Nurse Coordinator) for her assistance in recruiting volunteer subjects and coordinating sample collection. Additionally, Evan Linton from the CCTR provided critical assistance by designing and manufacturing the flexible plastic templates to enable reproducible replicate skin swab sampling. We would like to thank all of the members of the Lieberman Lab for valuable discussions and iterative protocol development which contributed to the methods described in this paper.

We would like to thank Bio-Me for informative discussions as well as their generous gift of qPCR panel analysis. We would like to thank technical and customer support staff at QIAGEN for useful discussions around the technology underlying the DNeasy 96 PowerSoil Pro kit and for providing us with materials to carry out PowerSoil Pro 96 DNA extractions.

Finally, we would like to thank the 20 people who volunteered as subjects for this study for their time and willingness to contribute samples to this study.

This work was funded by NIH grant 1DP2GM140922 to TDL.

