## Supplemental Figures 1-10 for "Microbiome diversity of low biomass skin sites is captured by metagenomics but not 16S amplicon sequencing"

Fig. S1

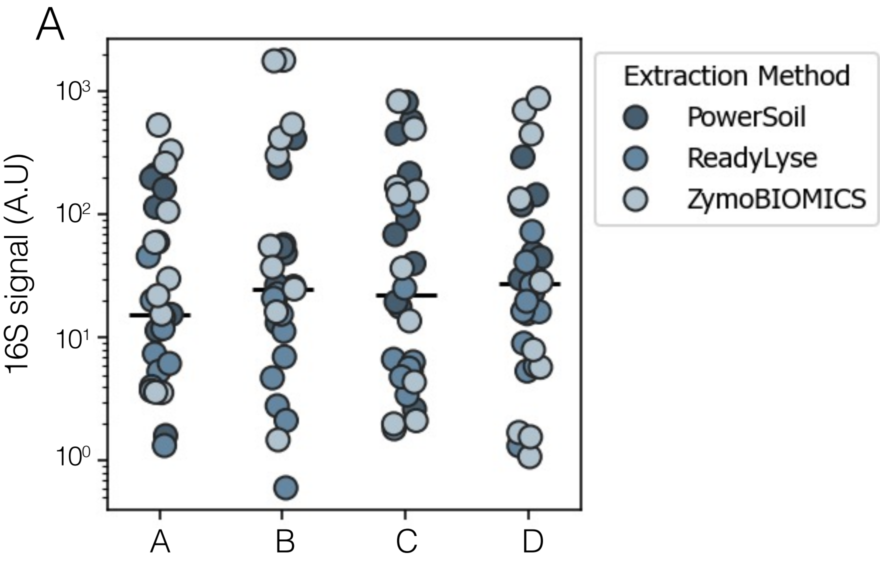


B

**Sample tubes:**

Tube 1: Zymogen BeadBashing Lysis Tube
Tube 2: 2ml tube with 1ml TESX
Tube 3: 0.5ml tube with 100ul TESX
Tube 4: 2ml tube with 1ml 50:50 glycerol:PBS

| Subject | A | B | C | D |
|---|---|---|---|---|
| 1 | Tube 1 | Tube 2 | Tube 3 | Tube 4 |
| 2 | Tube 2 | Tube 3 | Tube 4 | Tube 1 |
| 3 | Tube 3 | Tube 4 | Tube 1 | Tube 2 |
| 4 | Tube 4 | Tube 1 | Tube 2 | Tube 3 |
| … | Repeat tube order of first subject, then iterate through tube and quadrant assignments as above | | | |
| 20 | Tube 4 | Tube 1 | Tube 2 | Tube 3 |

**Fig. S1: Replicate skin swab samples were collected in randomized fashion – order of collection did not impact bacterial biomass**

**A)** There is no significant difference in bacterial biomass, as quantified by universal 16S qPCR, between the first sample collected (quadrant A) and the last sample collected (quadrant D) per subject per site, regardless of DNA extraction method used (indicated by color of symbol). Dots represent samples, bar indicates median. Quadrant: A N=29, B N=29, C N=30, D N=30. **B)** This panel describes the contents of each of the sample collection tubes used for downstream DNA extraction methods or preserved for future culture-based analysis (tube 4). All samples were stored at -80C° until DNA extraction. The table describes the randomization process used to determine which replicate sample quadrant of the forehead or leg was assigned to which collection tube.

Fig. S2

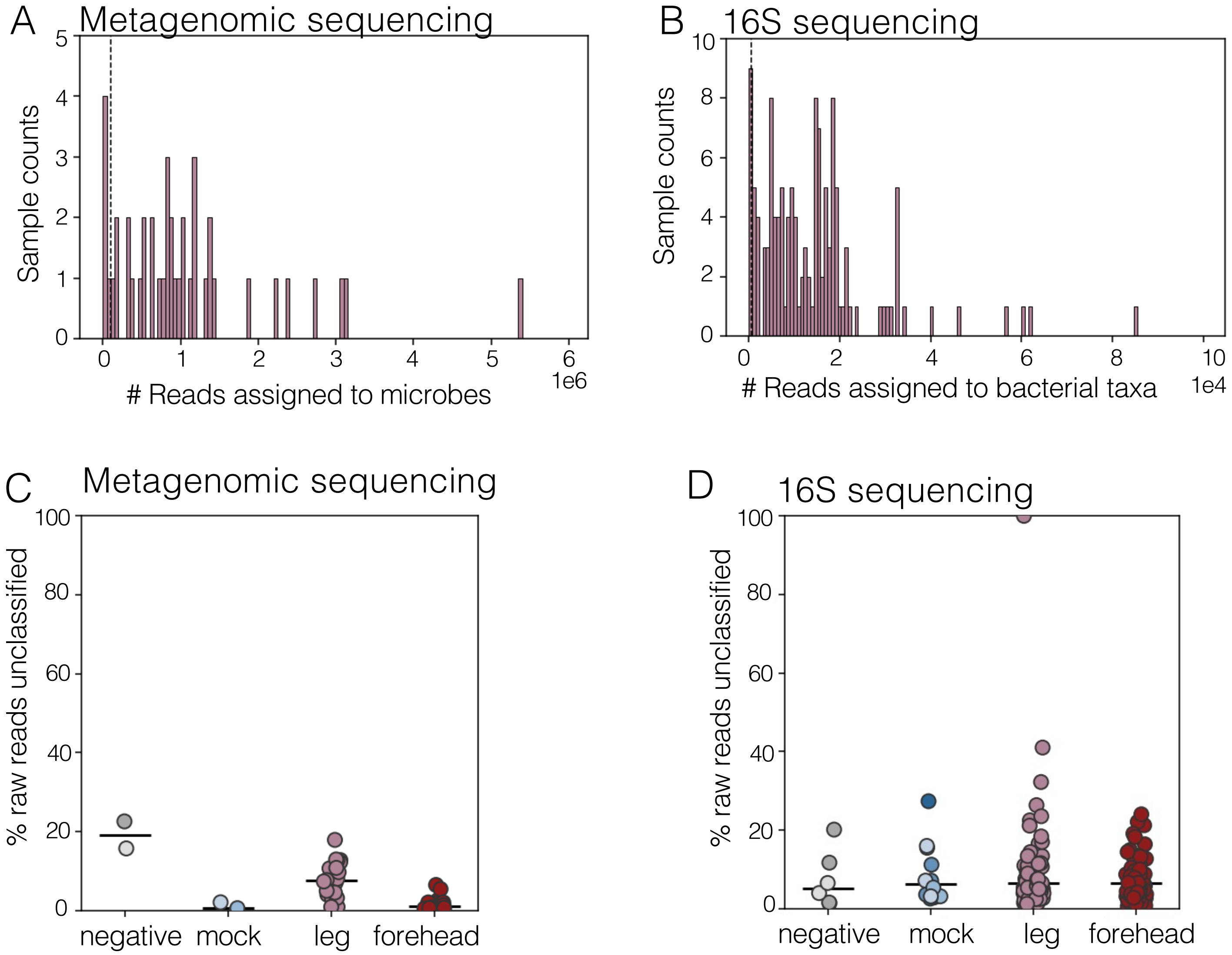


**Fig. S2: Majority of metagenomic reads were classified by kraken2 although variation was observed**
**A)** Metagenomic sequencing dataset: skin swab samples with fewer than 100,000 reads assigned to microbes by kraken2 were removed from analysis (5/20 leg samples removed, 0/20 forehead samples removed). All reads assigned to species classified as "Bacteria" or "Fungi" were included as microbial reads. **B)** 16S sequencing dataset: skin swab samples with fewer than 500 reads assigned to bacterial taxa (assigned a lower taxonomic level than "Bacteria") were discarded (6/120 leg samples and 2/120 forehead samples removed). **C)** The percentage of nonhuman reads not assigned to a microbial taxon in the metagenomics dataset is shown. Negative control samples had the highest unclassified read percentage likely reflecting the low number of total reads in these samples. As expected, mock community samples composed of five species had the lowest percentage of unclassified reads (0.1-2%, median of 0.55%). Forehead samples had a lower percentage of unclassified reads (0.14-6.4%, median of 1.1%) than leg samples (0.87-18%, median of 7.7%), likely reflecting the decreased diversity and increased *Cutibacterium* dominance of forehead samples compared to lower leg samples. **D)** In contrast, for 16S sequencing all sample types show a similar median unclassified reads of ~5-6%. This likely reflects the high proportion of *Cutibacterium* found in all 16S sequencing microbiome profiles across samples and negative controls (Fig. 2, Fig. S4 and Fig. S8). Metagenomics: leg N=13 and forehead N=20. 16S: leg N=58, forehead N=60.

Fig. S3

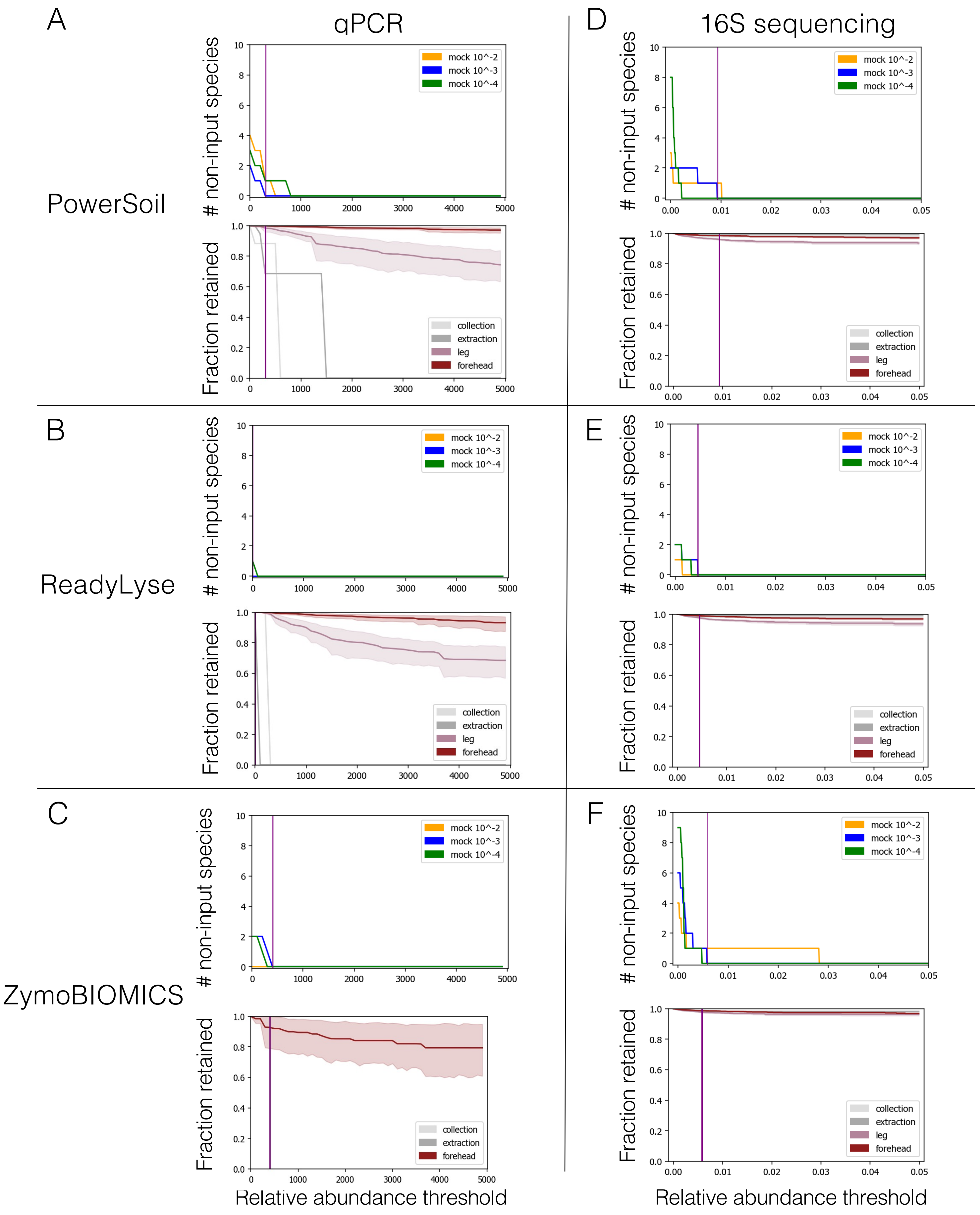
A
qPCR
D
16S sequencing
PowerSoil
B
ReadyLyse
E
C
ZymoBIOMICS
F
### non-input species
Fraction retained
mock 10^-2
mock 10^-3
mock 10^-4
collection
extraction
leg
forehead
Relative abundance threshold
Relative abundance threshold

**Fig. S3: Choosing a threshold for taxa filtering for qPCR and 16S datasets**
A threshold for removing likely contaminant taxa was chosen for each dataset -- unique combination of DNA extraction method and DNA analysis method. The top panel in each subplot shows how the threshold was chosen while the bottom panel shows the fraction of the original sample retained at each possible threshold. See Fig. 2 and Methods for additional details.

**Left (A-C):** Samples extracted with PS and RL and analyzed by qPCR show that higher biomass forehead samples retain a higher fraction of starting composition than low biomass leg skin samples, as expected due to the increased relative abundance of contaminant taxa in low biomass samples. Forehead samples extracted with the ZY kit analyzed by qPCR panel appear comparable to forehead samples in other qPCR datasets (no ZY leg samples included in analysis due to loss due to evaporation prior to qPCR panel analysis). There was no signal detected by qPCR in the ZY negative control samples thus no gray lines appear in the lower panel of C. qPCR analysis PS: leg N=18, forehead N=20. RL: leg N=20, forehead N=20. ZY: forehead N=14.

**Right (D-F):** 16S datasets show minimal changes in response to taxa filtering across forehead, leg and negative control samples. This reflects the high proportion of *Cutibacterium* in all 16S datasets across sample types– because very few low abundance taxa were detected in any sample, there was very little change when a threshold relative abundance filter was used to eliminate taxa below a certain relative abundance threshold. 16S sequencing analysis PS: leg N=18, forehead N=20. RL: leg N=17, forehead N=18. ZY: leg N=18, forehead N=20.

Fig. S4

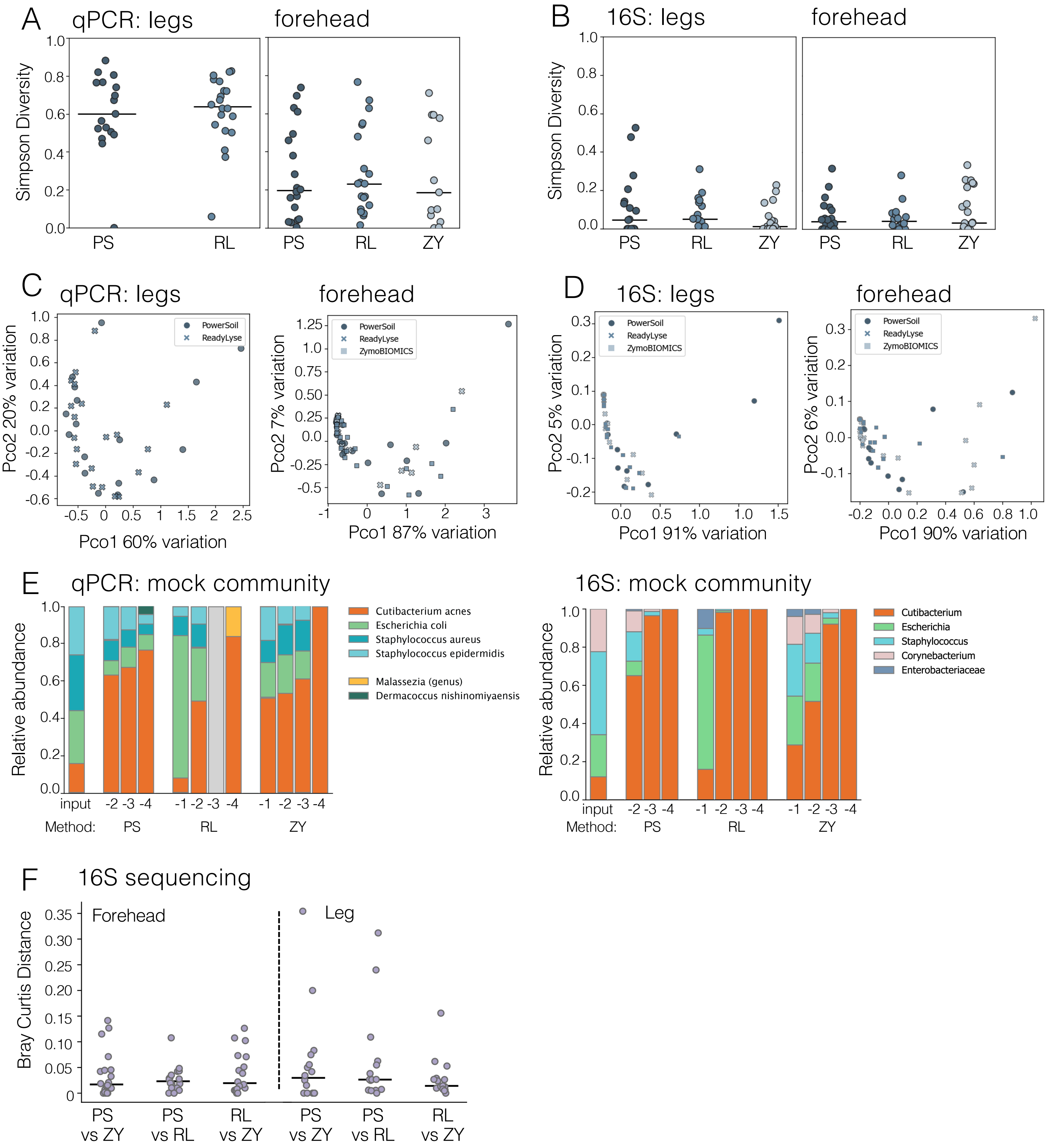
A
qPCR: legs
forehead
Simpson Diversity
1.0
0.8
0.6
0.4
0.2
0.0
PS
RL
ZY
B
16S: legs
C
Pco2 20% variation
Pco1 60% variation
PowerSoil
ReadyLyse
ZymoBIOMICS
Pco2 7% variation
Pco1 87% variation
D
Pco2 5% variation
Pco1 91% variation
Pco2 6% variation
Pco1 90% variation
E
qPCR: mock community
Relative abundance
Cutibacterium acnes
Escherichia coli
Staphylococcus aureus
Staphylococcus epidermidis
Malassezia (genus)
Dermacoccus nishinomiyaensis
input
Method:
16S: mock community
Cutibacterium
Escherichia
Staphylococcus
Corynebacterium
Enterobacteriaceae
F
16S sequencing
Bray Curtis Distance
Forehead
Leg
PS vs ZY
PS vs RL
RL vs ZY

**Fig. S4: DNA extraction method does not significantly change microbiome composition**
**A-B)** Both qPCR **(A)** and 16S sequencing **(B)** showed that there was no difference in diversity (Simpson metric) as a consequence of DNA extraction method for either low biomass leg samples (left) or high biomass forehead samples (right). **C-D)** Principal coordinates analysis using either qPCR (C) or 16S sequencing (D) showed that there was no significant separation between samples based on DNA extraction methods for leg samples (PERMANOVA P=0.45 and P=0.23) or forehead samples (PERMANOVA P=0.43 and P=0.35). For A-D, dots indicate samples and bars indicate median. **E)** The composition of mock community dilutions varied based on both extraction and analysis method. By qPCR we observe that PS is the most successful method, as it retains all four input species at the $10^{-4}$ dilution. However, by 16S sequencing the $10^{-4}$ dilution of both PS and ZY are exclusively *Cutibacterium* and the $10^{-3}$ dilution of ZY retains more input genera than does PS. By either analysis method, RL has the worst recall as the mock community is diluted. This is consistent with our recommendation based on sample yield that either kit-based method with combined mechanical and chemical lysis is preferable to the enzyme-only lysis of the RL method. **F)** In line in qPCR panel analysis (Fig. 4B), by 16S sequencing there were no significant differences between the Bray-Curtis distance separating replicate sample pairs between any of the DNA extraction method ($P>0.4$ for all). Forehead: PS N=20, RL N=18, ZY N=20. Leg: PS N=18, RL N=17, ZY N=18. Dots indicate sample pairs and lines indicate median. Significant p-values shown in bold.

Fig. S5

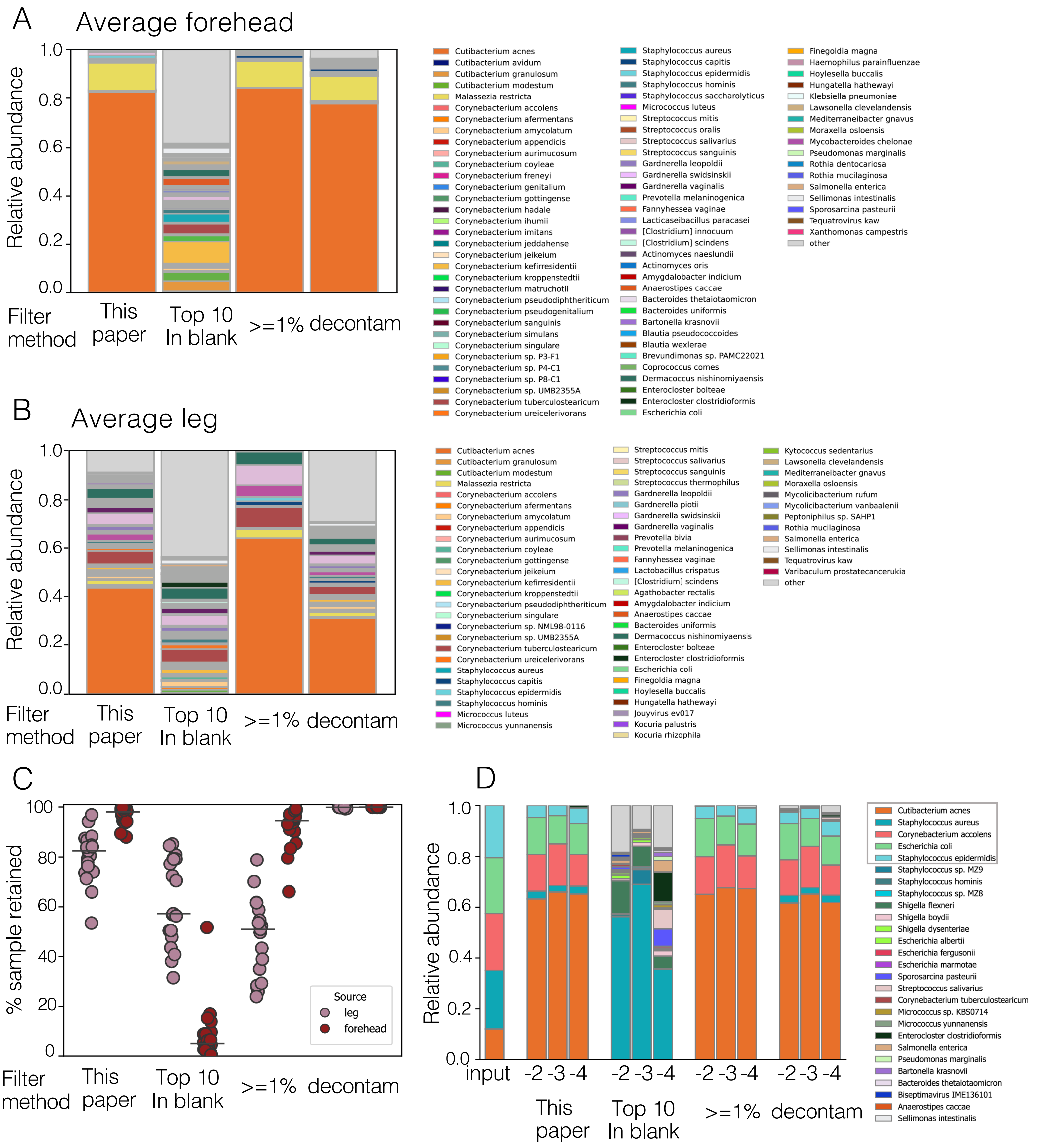
A
Average forehead
Relative abundance
1.0
0.8
0.6
0.4
0.2
0.0
Filter method
This paper
Top 10 In blank
>=1%
decontam
Cutibacterium acnes
Cutibacterium avidum
Cutibacterium granulosum
Cutibacterium modestum
Malassezia restricta
Corynebacterium accolens
Corynebacterium afermentans
Corynebacterium amycolatum
Corynebacterium appendicis
Corynebacterium aurimucosum
Corynebacterium coyleae
Corynebacterium freneyi
Corynebacterium genitalium
Corynebacterium gottingense
Corynebacterium hadale
Corynebacterium ihumii
Corynebacterium imitans
Corynebacterium jeddahense
Corynebacterium jeikeium
Corynebacterium kefirresidentii
Corynebacterium kroppenstedtii
Corynebacterium matruchotii
Corynebacterium pseudodiphtheriticum
Corynebacterium pseudogenitalium
Corynebacterium sanguinis
Corynebacterium simulans
Corynebacterium singulare
Corynebacterium sp. P3-F1
Corynebacterium sp. P4-C1
Corynebacterium sp. P8-C1
Corynebacterium sp. UMB2355A
Corynebacterium tuberculostearicum
Corynebacterium ureicelerivorans
Staphylococcus aureus
Staphylococcus capitis
Staphylococcus epidermidis
Staphylococcus hominis
Staphylococcus saccharolyticus
Micrococcus luteus
Streptococcus mitis
Streptococcus oralis
Streptococcus salivarius
Streptococcus sanguinis
Gardnerella leopoldii
Gardnerella swidsinskii
Gardnerella vaginalis
Prevotella melaninogenica
Fannyhessea vaginae
Lacticaseibacillus paracasei
[Clostridium] innocuum
[Clostridium] scindens
Actinomyces naeslundii
Actinomyces oris
Amygdalobacter indicium
Anaerostipes caccae
Bacteroides thetaiotaomicron
Bacteroides uniformis
Bartonella krasnovii
Blautia pseudococcoides
Blautia wexlerae
Brevundimonas sp. PAMC22021
Coprococcus comes
Dermacoccus nishinomiyaensis
Enterocloster bolteae
Enterocloster clostridioformis
Escherichia coli
Finegoldia magna
Haemophilus parainfluenzae
Hoylesella buccalis
Hungatella hathewayi
Klebsiella pneumoniae
Lawsonella clevelandensis
Mediterraneibacter gnavus
Moraxella osloensis
Mycobacteroides chelonae
Pseudomonas marginalis
Rothia dentocariosa
Rothia mucilaginosa
Salmonella enterica
Sellimonas intestinalis
Sporosarcina pasteurii
Tequatrovirus kaw
Xanthomonas campestris
other
B
Average leg
Relative abundance
1.0
0.8
0.6
0.4
0.2
0.0
Filter method
This paper
Top 10 In blank
>=1%
decontam
Cutibacterium acnes
Cutibacterium granulosum
Cutibacterium modestum
Malassezia restricta
Corynebacterium accolens
Corynebacterium afermentans
Corynebacterium amycolatum
Corynebacterium appendicis
Corynebacterium aurimucosum
Corynebacterium coyleae
Corynebacterium gottingense
Corynebacterium jeikeium
Corynebacterium kefirresidentii
Corynebacterium kroppenstedtii
Corynebacterium pseudodiphtheriticum
Corynebacterium singulare
Corynebacterium sp. NML98-0116
Corynebacterium sp. UMB2355A
Corynebacterium tuberculostearicum
Corynebacterium ureicelerivorans
Staphylococcus aureus
Staphylococcus capitis
Staphylococcus epidermidis
Staphylococcus hominis
Micrococcus luteus
Micrococcus yunnanensis
Streptococcus mitis
Streptococcus salivarius
Streptococcus sanguinis
Streptococcus thermophilus
Gardnerella leopoldii
Gardnerella piotii
Gardnerella swidsinskii
Gardnerella vaginalis
Prevotella bivia
Prevotella melaninogenica
Fannyhessea vaginae
Lactobacillus crispatus
[Clostridium] scindens
Agathobacter rectalis
Amygdalobacter indicium
Anaerostipes caccae
Bacteroides uniformis
Dermacoccus nishinomiyaensis
Enterocloster bolteae
Enterocloster clostridioformis
Escherichia coli
Finegoldia magna
Hoylesella buccalis
Hungatella hathewayi
Jouyvirus ev017
Kocuria palustris
Kocuria rhizophila
Kytococcus sedentarius
Lawsonella clevelandensis
Mediterraneibacter gnavus
Moraxella osloensis
Mycolicibacterium rufum
Mycolicibacterium vanbaalenii
Peptoniphilus sp. SAHP1
Rothia mucilaginosa
Salmonella enterica
Sellimonas intestinalis
Tequatrovirus kaw
Varibaculum prostatecancerukia
other
C
% sample retained
100
80
60
40
20
0
Source
leg
forehead
Filter method
This paper
Top 10 In blank
>=1%
decontam
D
Relative abundance
1.0
0.8
0.6
0.4
0.2
0.0
input
-2 -3 -4
-2 -3 -4
-2 -3 -4
-2 -3 -4
This paper
Top 10 In blank
>=1%
decontam
Cutibacterium acnes
Staphylococcus aureus
Corynebacterium accolens
Escherichia coli
Staphylococcus epidermidis
Staphylococcus sp. MZ9
Staphylococcus hominis
Staphylococcus sp. MZ8
Shigella flexneri
Shigella boydii
Shigella dysenteriae
Escherichia albertii
Escherichia fergusonii
Escherichia marmotae
Sporosarcina pasteurii
Streptococcus salivarius
Corynebacterium tuberculostearicum
Micrococcus sp. KBS0714
Micrococcus yunnanensis
Enterocloster clostridioformis
Salmonella enterica
Pseudomonas marginalis
Bartonella krasnovii
Bacteroides thetaiotaomicron
Biseptimavirus IME136101
Anaerostipes caccae
Sellimonas intestinalis

**Fig. S5: Different taxa filtering strategies alter swab and mock community composition**
This figure compares the taxa filtering method used in this paper to alternate published filtering methods using the metagenomic sequencing dataset. Alternate filtering methods include “Top 10 in blank”: removing the most abundant taxa present in the negative control samples, “>=1%”: removing all taxa with an average relative abundance across samples lower than a threshold value (here we chose 1%) and “decontam”: using the R package decontam to analyze the unfiltered dataset and removing those taxa it identified as likely contaminants.

**A)** The filtering method used in this paper, “>=1%” and decontam all result in a highly similar average forehead skin microbiome which is dominated by *Cutibacterium acnes* and *Malassezia restricta*, with a smaller relative abundance of other expected skin microbes including *Staphylococcus capitis* and *Staphylococcus epidermidis*. In contrast, the “top 10 in blank” method removes *C. acnes* and several *Staphylococcus* species such that the most abundant species is *Corynebacterium kefirresidentii*. This supports our conclusion that wholesale removal of taxa found in negative controls is not an appropriate filtering method. The 50 most abundant species on average are shown in color**. B)** For leg skin microbiome samples, again, the method used in this paper as well as “>=1%” and decontam accurately show that the most abundant skin microbe is *Cutibacterium acnes,* while the “top 10 in blank” method inaccurately removes this and many other common skin microbes. The “>=1%” method also removes all abundance species, while decontam retains more and the method used in this paper falls somewhere in the middle. The 50 most abundant species on average are shown in color. **C)** For the forehead samples, all of the filter methods except for “top 10 in blank” had a comparable and minimal impact on the percentage of sample retained. For the leg skin samples, decontam retained the most of the original sample, “top 10 in blank” and “>=1%” were comparable and retained the least, and the method used in this paper was in the middle. **D)** For mock community samples, the “top10 in blank” method fails to include four out of five input species and inaccurately retains many non-input species. Both the method used in this paper and the “>=1%” method successfully recall all five species and only have non-input signal present in the lowest dilution. The decontam method successfully recalls all five species and is closer to the input ratios but fails to filter non-input species from all three dilutions. Throughout, leg N=13 and forehead N=20.

Fig. S6

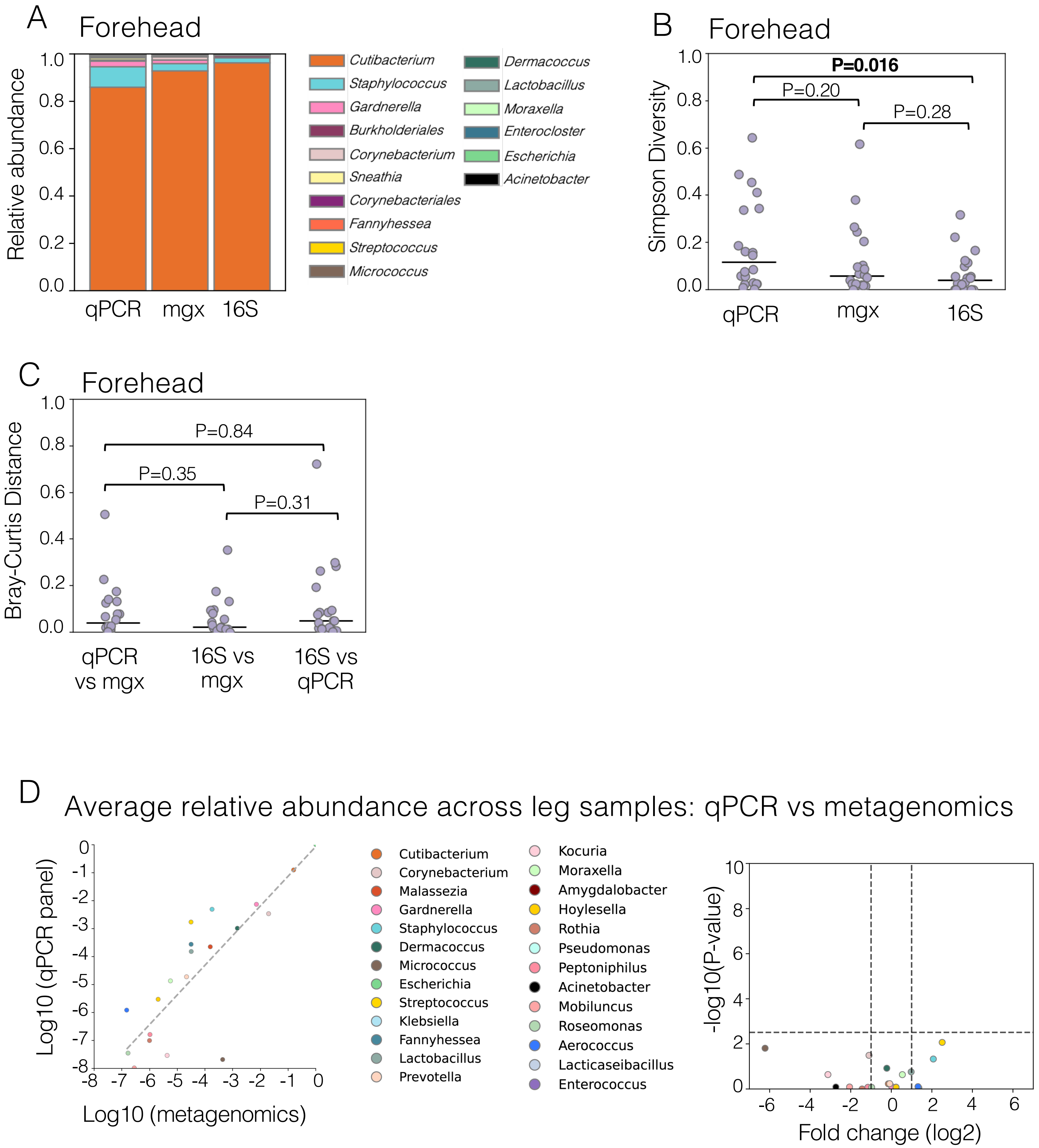
A
Forehead
Relative abundance
1.0
0.8
0.6
0.4
0.2
0.0
qPCR
mgx
16S
Cutibacterium
Staphylococcus
Gardnerella
Burkholderiales
Corynebacterium
Sneathia
Corynebacteriales
Fannyhessea
Streptococcus
Micrococcus
Dermacoccus
Lactobacillus
Moraxella
Enterocloster
Escherichia
Acinetobacter
B
Forehead
Simpson Diversity
P=0.016
P=0.20
P=0.28
qPCR
mgx
16S
C
Forehead
Bray-Curtis Distance
P=0.84
P=0.35
P=0.31
qPCR vs mgx
16S vs mgx
16S vs qPCR
D
Average relative abundance across leg samples: qPCR vs metagenomics
Log10 (qPCR panel)
Log10 (metagenomics)
Cutibacterium
Corynebacterium
Malassezia
Gardnerella
Staphylococcus
Dermacoccus
Micrococcus
Escherichia
Streptococcus
Klebsiella
Fannyhessea
Lactobacillus
Prevotella
Kocuria
Moraxella
Amygdalobacter
Hoylesella
Rothia
Pseudomonas
Peptoniphilus
Acinetobacter
Mobiluncus
Roseomonas
Aerococcus
Lacticaseibacillus
Enterococcus
-log10(P-value)
Fold change (log2)

**Fig. S6: DNA analysis by qPCR and metagenomics is highly concordant**

**A)** The same genera in largely similar proportions were detected regardless of DNA analysis method (qPCR panel, metagenomics or 16S) in the low diversity forehead skin microbiome samples. **B)** The per sample microbiome diversity (Simpson metric) of forehead skin samples was significantly higher when measured by qPCR panel compared to 16S sequencing (P=0.016) but qPCR and metagenomics were comparable (P=0.2). There was no significant difference between metagenomic and 16S sequencing, in contrast with the equivalent comparison across leg microbiome samples (Fig. 3) **C)** We measured the Bray-Curtis distance between the same sample analyzed by different DNA analysis methods and found that the difference between qPCR and metagenomics samples is comparable to the distance between qPCR and 16S sequencing samples as well as the distance between profiles generated by 16S and metagenomic sequencing. This suggests that while the individual sample diversity was higher in the qPCR dataset, the composition of samples based on shared genera was highly similar between analysis methods**. D)** We compared the average relative abundance of microbial genera as measured by qPCR or metagenomics and observed a high correlation in relative abundance (Pearson correlation, $R^2$=0.90, P=$4.23 \times 10^{-11}$) and that no genus was significantly different in relative abundance between methods. Fold-change (right) refers to qPCR relative abundance divided by metagenomics relative abundance. Forehead samples: N=20 for all DNA analysis methods. Leg samples (D): N=13 metagenomics and N=18 qPCR panel.

Fig. S7

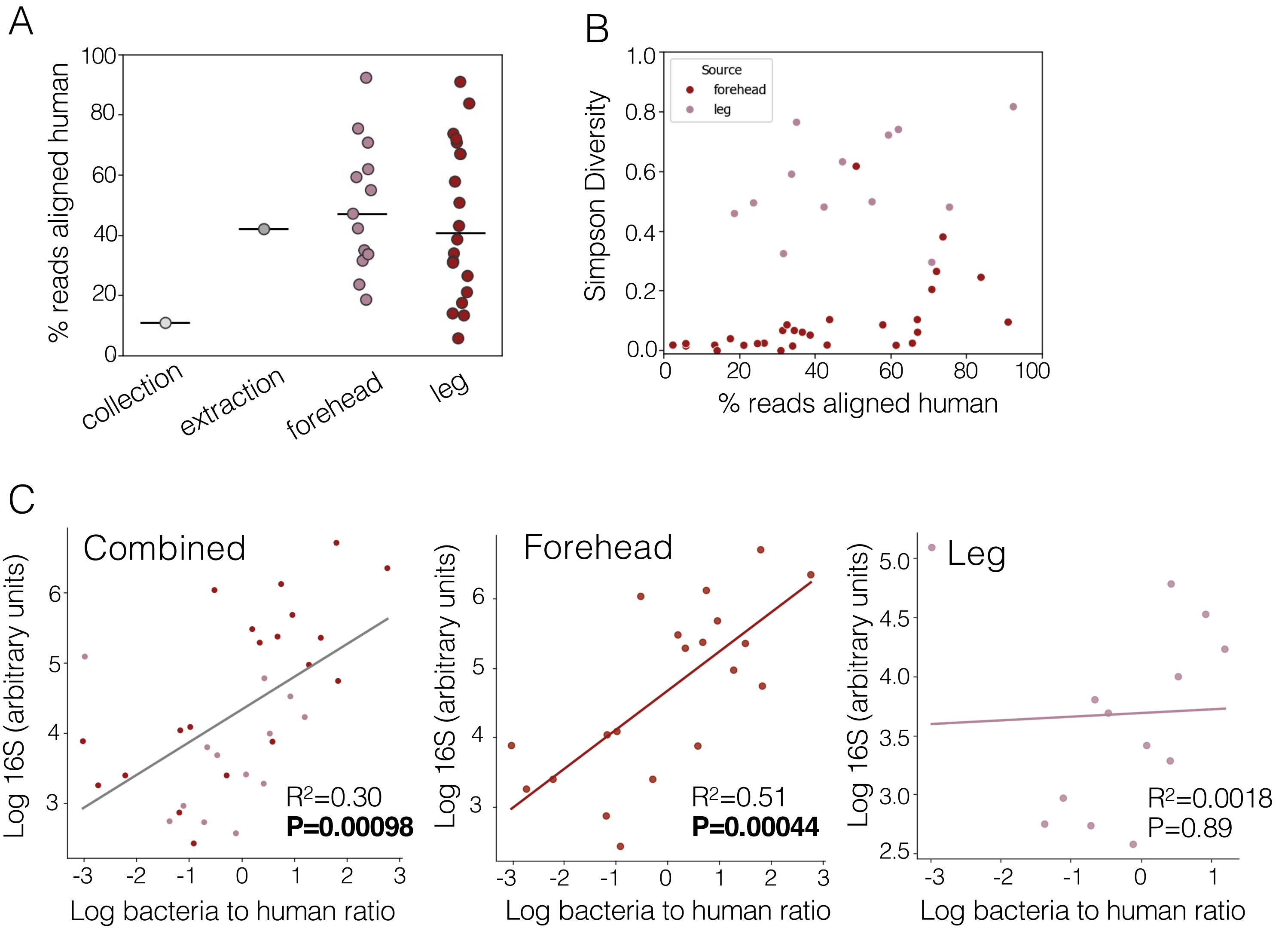


**Fig. S7: Human reads present in metagenomic samples do not limit microbiome diversity and can be used to infer total microbial DNA content in high biomass skin swab samples**

**A)** Skin swab samples from both the high microbial biomass forehead and the low microbial biomass leg skin show a wide range of percent sample human reads. One of the two negative controls is comparable to the median leg sample, suggesting contamination of this sample from a skin swab sample, in line with the composition of this control (Fig. 2A). Lines indicate median. **B)** The percentage of human reads per sample does not correlate with microbiome diversity in either forehead or leg skin swabs, demonstrating that the presence of human DNA in the skin swab sample does not inhibit profiling of microbial diversity. **C)** We observe a significant correlation between bacterial to human read ratio and 16S copy number when assessing all skin swab samples collectively ($R^2$=0.30, P=0.00098) or the high biomass forehead skin samples alone ($R^2$=0.51, P=0.00044) but no correlation ($R^2$=0.0018, P=0.89) in the low biomass leg skin samples, suggesting that this mode of normalization may only be appropriate for high microbial biomass sites. Bacterial reads were those assigned to "Bacteria" taxa by kraken2 and human reads are those aligned to human genome by bowtie2. Dots indicate samples and lines indicate the Pearson correlation between values. Throughout, forehead N=20 and leg N=13.

Fig. S8

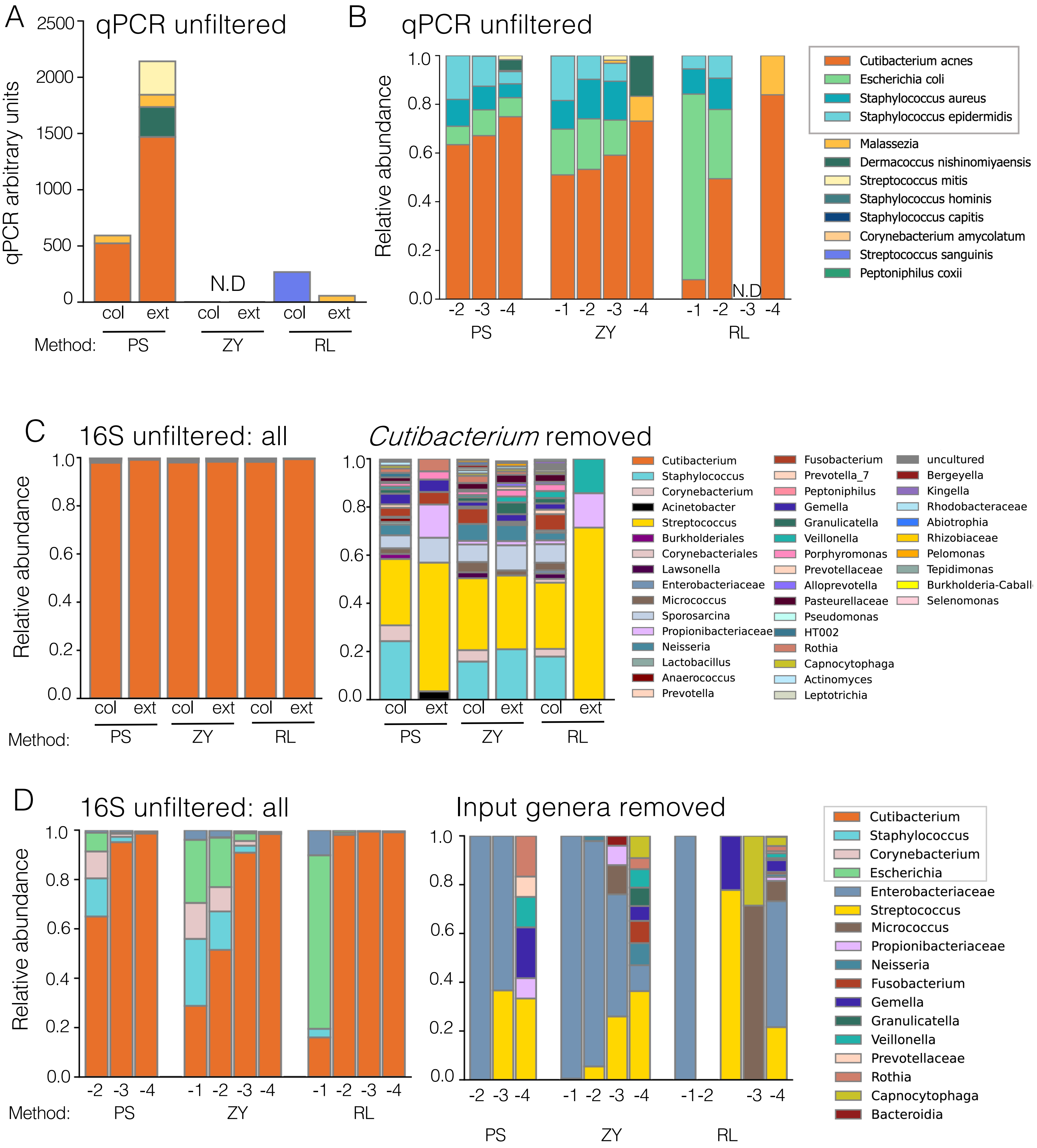

A
qPCR unfiltered
qPCR arbitrary units
2500
2000
1500
1000
500
0
N.D
col ext col ext col ext
Method: PS ZY RL
B
qPCR unfiltered
Relative abundance
1.0
0.8
0.6
0.4
0.2
0.0
N.D
-2 -3 -4 -1 -2 -3 -4 -1 -2 -3 -4
PS ZY RL
Cutibacterium acnes
Escherichia coli
Staphylococcus aureus
Staphylococcus epidermidis
Malassezia
Dermacoccus nishinomiyaensis
Streptococcus mitis
Staphylococcus hominis
Staphylococcus capitis
Corynebacterium amycolatum
Streptococcus sanguinis
Peptoniphilus coxii
C
16S unfiltered: all
Cutibacterium removed
Relative abundance
col ext col ext col ext
Method: PS ZY RL
Cutibacterium
Staphylococcus
Corynebacterium
Acinetobacter
Streptococcus
Burkholderiales
Corynebacteriales
Lawsonella
Enterobacteriaceae
Micrococcus
Sporosarcina
Propionibacteriaceae
Neisseria
Lactobacillus
Anaerococcus
Prevotella
Fusobacterium
Prevotella_7
Peptoniphilus
Gemella
Granulicatella
Veillonella
Porphyromonas
Prevotellaceae
Alloprevotella
Pasteurellaceae
Pseudomonas
HT002
Rothia
Capnocytophaga
Actinomyces
Leptotrichia
uncultured
Bergeyella
Kingella
Rhodobacteraceae
Abiotrophia
Rhizobiaceae
Pelomonas
Tepidimonas
Burkholderia-Caball
Selenomonas
D
16S unfiltered: all
Input genera removed
Relative abundance
-2 -3 -4 -1 -2 -3 -4 -1 -2 -3 -4
Method: PS ZY RL
-2 -3 -4 -1 -2 -3 -4 -1 -2 -3 -4
PS ZY RL
Cutibacterium
Staphylococcus
Corynebacterium
Escherichia
Enterobacteriaceae
Streptococcus
Micrococcus
Propionibacteriaceae
Neisseria
Fusobacterium
Gemella
Granulicatella
Veillonella
Prevotellaceae
Rothia
Capnocytophaga
Bacteroidia

**Fig. S8: Composition of negative controls is largely consistent across DNA extraction methods and differs between DNA analysis methods.**

**A)** There was increased absolute abundance of bacterial DNA in the PS negative control samples compared to either the ZY or RL samples, in line with other observations that the PS lysis plate increased likelihood of well-to-well transfer of bacterial DNA. **B)** By qPCR, PS and ZY mock community samples have a more consistent composition across the dilution series than RL samples. However, across all DNA extraction methods, non-input species become more abundant as the mock community sample is diluted. Input species shown in the gray box. **C)** Across extraction methods, the negative controls are >90% *Cutibacterium* by relative abundance. When *Cutibacterium* is removed from analysis (right), we observed that the extraction controls from PS and RL have notably fewer non-*Cutibacterium* contaminants than other negative control samples but still share many contaminants with ZY and collection negative controls. We conclude that the dominant source of contaminant DNA was well-to-well contamination from surrounding skin swab samples rather than method-specific reagents. **D)** Analysis of the total unfiltered mock community samples (left) and the non-input genera only (right) similarly shows multiple contaminant bacterial genera across DNA extraction methods and no strong association between DNA extraction method and any single genus.

Fig. S9

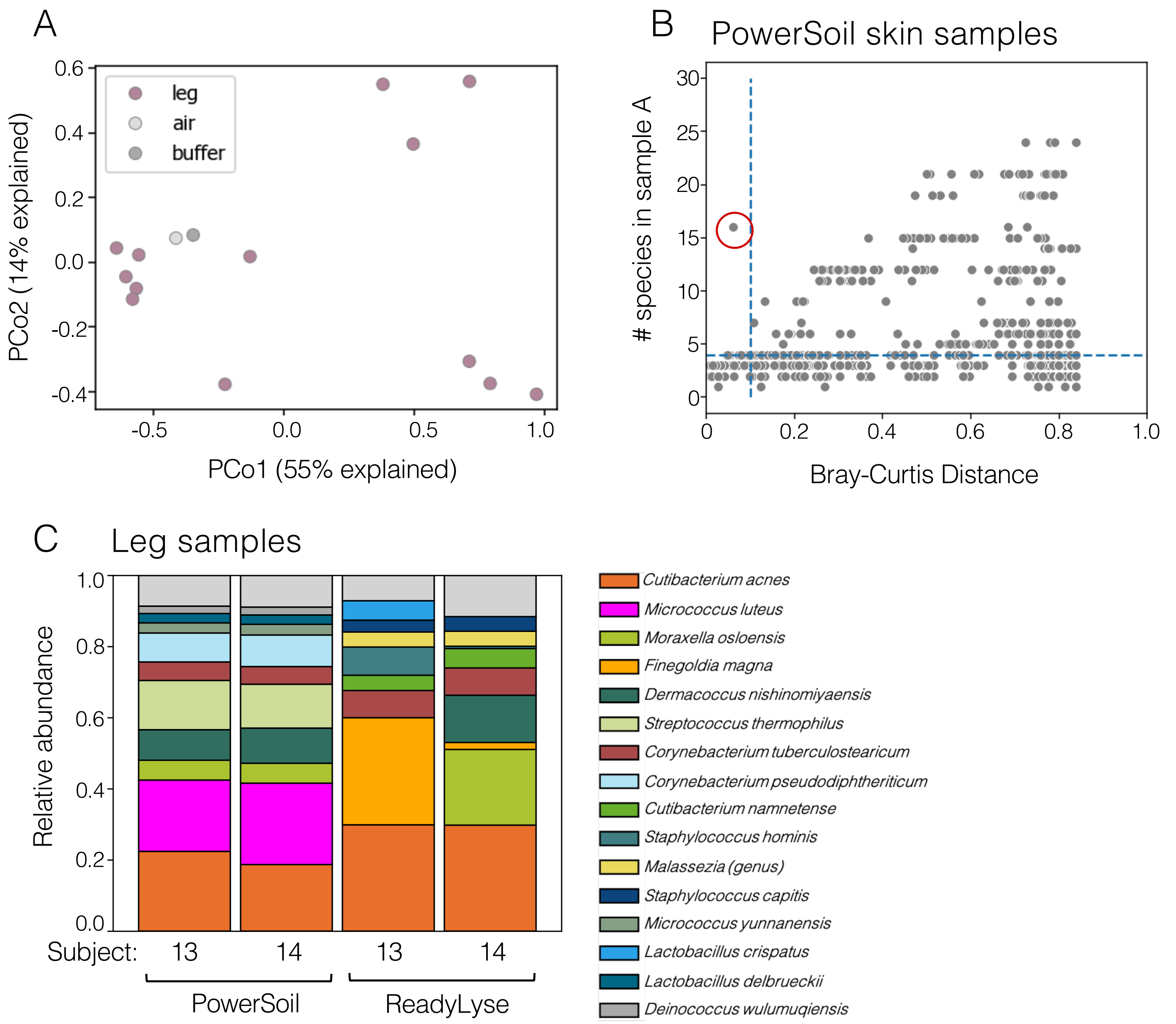


**Fig. S9: The Bray-Curtis distance between samples was used to determine whether leg samples were distinct from controls and identify well-to-well contamination**

**A)** After sample and taxa filtering, we used principal coordinates analysis to visualize the Bray-Curtis distance between leg skin samples and negative controls and observed separation between low biomass leg skin samples and negative controls (PERMANOVA P=0.15), supporting the conclusion that this workflow is sufficient to enable accurate analysis of low biomass leg microbiome samples. Leg N=13. **B)** To identify samples potentially contaminated by neighboring wells, we measured the Bray-Curtis distance between samples from different subjects within a DNA extraction method. The pair of samples circled in red has a Bray-Curtis distance <0.1 and more than five species present per sample, highlighting it as a possible contaminated pair. **C)** For suspect pairs of samples, we compared the composition of the sample pairs by relative abundance across multiple DNA extraction methods. This plot shows the pair of samples circled in red in B, leg samples from subjects 13 and 14, which were highly similar in composition when extracted by the PowerSoil method, but which are not similar when extracted by the ReadyLyse method. This inconsistency suggests that the samples were contaminated during PowerSoil extraction and thus the sample in the pair with the lower biomass was discarded.

### Fig. S10

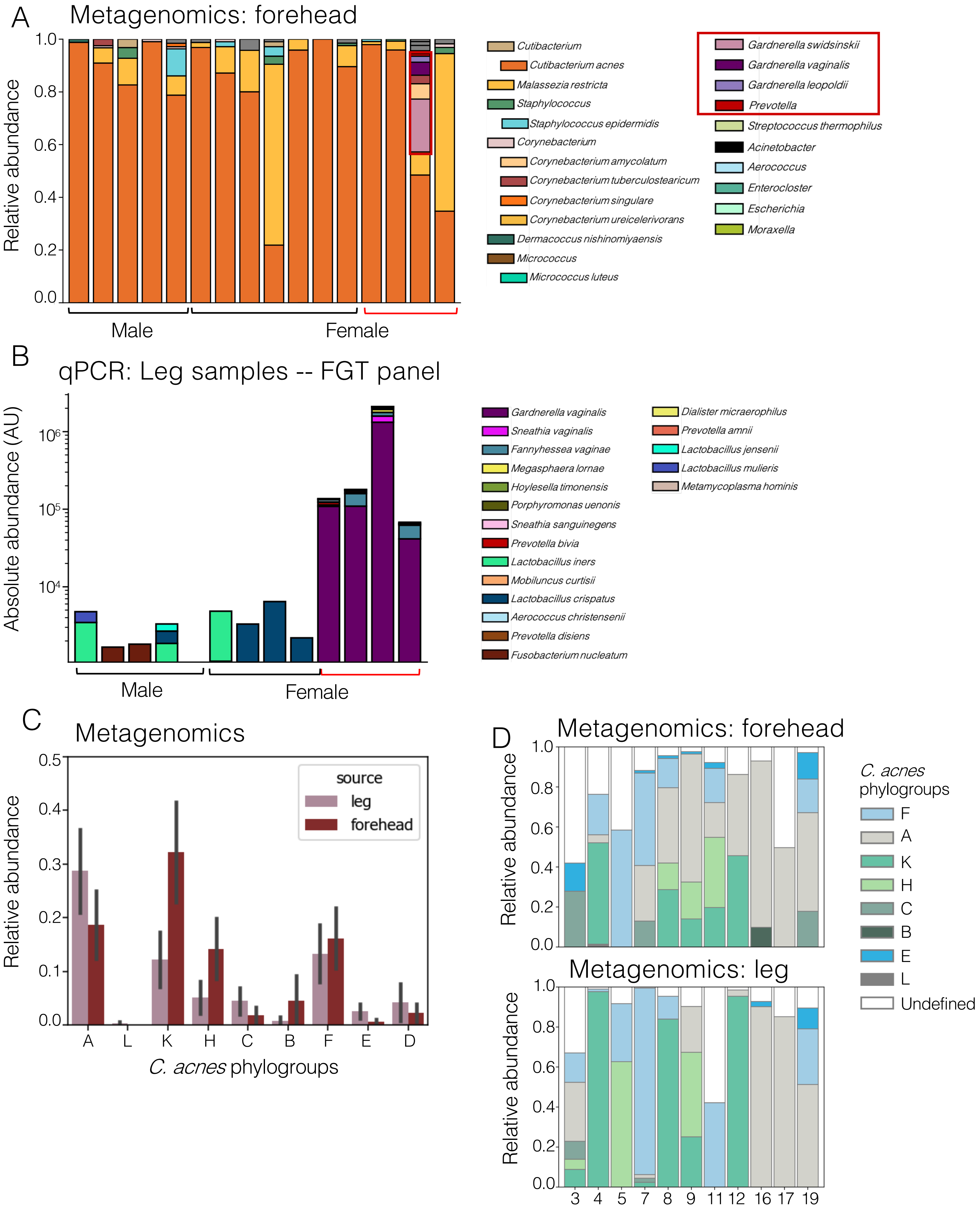
A
Metagenomics: forehead
Relative abundance
Male
Female
Cutibacterium
Cutibacterium acnes
Malassezia restricta
Staphylococcus
Staphylococcus epidermidis
Corynebacterium
Corynebacterium amycolatum
Corynebacterium tuberculostearicum
Corynebacterium singulare
Corynebacterium ureicelerivorans
Dermacoccus nishinomiyaensis
Micrococcus
Micrococcus luteus
Gardnerella swidsinskii
Gardnerella vaginalis
Gardnerella leopoldii
Prevotella
Streptococcus thermophilus
Acinetobacter
Aerococcus
Enterocloster
Escherichia
Moraxella
B
qPCR: Leg samples -- FGT panel
Absolute abundance (AU)
Male
Female
Gardnerella vaginalis
Sneathia vaginalis
Fannyhessea vaginae
Megasphaera lornae
Hoylesella timonensis
Porphyromonas uenonis
Sneathia sanguinegens
Prevotella bivia
Lactobacillus iners
Mobiluncus curtisii
Lactobacillus crispatus
Aerococcus christensenii
Prevotella disiens
Fusobacterium nucleatum
Dialister micraerophilus
Prevotella amnii
Lactobacillus jensenii
Lactobacillus mulieris
Metamycoplasma hominis
C
Metagenomics
Relative abundance
source
leg
forehead
C. acnes phylogroups
D
Metagenomics: forehead
Metagenomics: leg
Relative abundance
C. acnes phylogroups
F
A
K
H
C
B
E
L
Undefined

**Fig. S10: Metagenomics and qPCR analysis of microbiome samples shows species and strain-level sharing between forehead and lower leg skin microbiome sites**

**A)** Metagenomic sequencing analysis of the forehead skin microbiome is highly consistent across subjects and is dominated by *C. acnes* with the exception of two subjects, dominated by the fungus, *Malassezia restricta.* Of the four samples from subjects with leg samples dominated by FGT genera, one forehead sample is also dominated by the FGT genus *Gardnerella*. Species and genera present at a relative abundance of >=10% in at least one sample were given a unique color. FGT species enclosed in red. Samples corresponding to female subjects with >20% FGT species in leg skin swab samples are bracketed in red. **B)** An FGT microbiome qPCR panel shows that the four leg swab samples with high relative abundance of vaginal microbiome species by metagenomics (Fig. 5A) are also the only subjects with detectable *Gardnerella vaginalis* DNA by qPCR. All other subjects, male and female, have ~10-fold less total absolute abundance of vaginal microbiome species, largely from genera also found in the gut microbiome (*Lactobacillus* and *Fusobacterium*) in keeping with a lack of transfer from the FGT in these subjects. **C)** Sub-species relative abundance analysis (PHLAME) of *C. acnes* using the metagenomics dataset showed no significant differences in relative abundance for any phylogroup between the forehead and leg skin samples, supporting previous findings that individuals carry consistent populations of *C. acnes* across skin sites. Bars represent average phylogroup abundance and lines indicate the standard error of the mean. **D)** Barplots show the relative abundance of different *C. acnes* phylogroups per sample and highlight the similar ratios of phylogroups present on paired forehead (top) and leg (bottom) samples taken from the same subject. Only those subjects with a leg sample with >=25% defined *C. acnes* phylogroups were included in this plot (N=11 subjects). For A, B and D, bars represent individual samples. For metagenomics analysis (A and C), leg N=13, forehead N=20 and negative controls N=2. For qPCR analysis (B) leg N=18.
