## Supplementary material for "Microbiome diversity of low biomass skin sites is captured by metagenomics but not 16S amplicon sequencing": Table S11-S12

Bio-Me
Microbiome Profiling

### Core Vaginal Microbiome Test - Taxa List

| 47 bacterial taxa | |
|---|---|
| *Aerococcus christensenii* | *Megasphaera hutchinsoni (Megasphaera type 2)* |
| *Amygdalobacter indicium (BVAB2)* | *Megasphaera lornae (Megasphaera type 1)* |
| *Amygdalobacter nucleatus (BVAB2)* | *Megasphaera vaginalis* |
| *Bifidobacterium bifidum* | *Metamycoplasma hominis* |
| *Bifidobacterium breve* | *Mobiluncus curtisii* |
| *Bifidobacterium longum* | *Mobiluncus mulieris* |
| *Clostridiales genomosp. BVAB1* | *Porphyromonas asaccharolytica* |
| *Dialister micraerophilus* | *Porphyromonas uenonis* |
| *Enterococcus faecalis* | *Prevotella amnii* |
| *Enterococcus faecium* | *Prevotella bivia* |
| *Escherichia coli* | *Prevotella disiens* |
| *Fannyhessea vaginae* | *Sneathia sanguinegens* |
| *Finegoldia magna* | *Sneathia vaginalis* |
| *Fusobacterium nucleatum* | *Staphylococcus aureus* |
| *Gardnerella vaginalis* | *Streptococcus agalactiae* |
| *Hoylesella timonensis* | *Streptococcus anginosus* |
| *Klebsiella pneumoniae* | *Ureaplasma parvum* |
| *Lacticaseibacillus rhamnosus* | *Ureaplasma urealyticum* |
| *Lactobacillus crispatus* | **6 fungi taxa** |
| *Lactobacillus gasseri* | *Candida albicans* |
| *Lactobacillus helveticus* | *Candida dubliniensis* |
| *Lactobacillus iners* | *Candida parapsilosis* |
| *Lactobacillus jensenii* | *Candida tropicalis* |
| *Lactobacillus mulieris* | *Pichia kudriavzevii* |
| *Lactobacillus paragasseri* | *Nakaseomyces glabratus* |
| *Ligilactobacillus salivarius* | **3 controls** |
| *Limosilactobacillus fermentum* | 16S bacteria control |
| *Limosilactobacillus reuteri* | Human RNase P |
| *Mageeibacillus indolicus (BVAB3)* | Internal PCR control |

Bio-Me
Microbiome Profiling

### Skin microbiome panel v2

| 48 bacterial taxa | |
|---|---|
| *Acinetobacter baumannii* | *Peptoniphilus coxii* |
| *Acinetobacter johnsonii* | *Peptoniphilus lacrimalis* |
| *Acinetobacter lwoffii* | *Pseudomonas aeruginosa* |
| *Anaerococcus nagyae* | *Roseomonas mucosa* |
| *Anaerococcus octavius* | *Rothia dentocariosa* |
| *Anaerococcus vaginalis* | *Staphylococcus aureus* |
| *Corynebacterium amycolatum* | *Staphylococcus capitis* |
| *Corynebacterium bovis* | *Staphylococcus caprae* |
| *Corynebacterium jeikeium* | *Staphylococcus epidermidis* |
| *Corynebacterium kroppenstedtii* | *Staphylococcus haemolyticus* |
| *Corynebacterium matruchotii* | *Staphylococcus hominis* |
| *Corynebacterium simulans* | *Staphylococcus lugdunensis* |
| *Corynebacterium tuberculostearicum* | *Staphylococcus saprophyticus* |
| *Cutibacterium acnes* | *Staphylococcus warneri* |
| *Cutibacterium avidum* | *Streptococcus infantis ATCC 700779* |
| *Cutibacterium granulosum* | *Streptococcus mitis* |
| *Cutibacterium namnetense* | *Streptococcus salivarius* |
| *Dermacoccus nishinomiyaensis* | *Streptococcus sanguinis* |
| *Dietzia cinnamea* | *Streptococcus thermophilus* |
| *Dietzia maris* | 5 fungi taxa |
| *Escherichia coli* | *Malassezia arunalokei* |
| *Finegoldia magna* | *Malassezia furfur* |
| *Kocuria rhizophila* | *Malassezia globosa* |
| *Lactobacillus crispatus* | *Malassezia restricta* |
| *Lactobacillus iners* | *Malassezia sympodialis* |
| *Lactococcus lactis* | 3 controls |
| *Micrococcus luteus ATCC 12698* | 16S rRNA control |
| *Moraxella osloensis* | Total *Malassezia* control |
| *Nitrosomonas eutropha* | Synthetic DNA PCR control |

October 2023
